# Identification of a genetic element required for spore killing in Neurospora

**DOI:** 10.1101/404004

**Authors:** Nicholas A. Rhoades, Austin M. Harvey, Dilini A. Samarajeewa, Jesper Svedberg, Aykhan Yusifov, Anna Abusharekh, Pennapa Manitchotpisit, Daren W. Brown, Kevin J. Sharp, David G. Rehard, Joshua Peters, Xavier Ostolaza-Maldonado, Jackson Stephenson, Patrick K. T. Shiu, Hanna Johannesson, Thomas M. Hammond

**Author notes:** These authors contributed equally to this work (Austin M. Harvey, Dilini A. Samarajeewa). Corresponding author: 346 Science Laboratory Building, Illinois State University, School of Biological Sciences, Normal, IL 61790.

## Abstract

Meiotic drive elements like *Spore killer-2* (*Sk-2*) in Neurospora are transmitted through sexual reproduction to the next generation in a biased manner. *Sk-2* achieves this biased transmission through spore killing. Here, we identify *rfk-1* as a gene required for the spore killing mechanism. The *rfk-1* gene is associated with a 1,481 bp DNA interval (called *AH36*) near the right border of the 30 cM *Sk-2* element, and its deletion eliminates the ability of *Sk-2* to kill spores. The *rfk-1* gene also appears to be sufficient for spore killing because its insertion into a non-*Sk-2* isolate disrupts sexual reproduction after the initiation of meiosis. Although the complete *rfk-1* transcript has yet to be defined, our data indicate that *rfk-1* encodes a protein of at least 39 amino acids and that *rfk-1* has evolved from a partial duplication of gene *ncu07086*. We also present evidence that *rfk-1*’s location near the right border of *Sk-2* is critical for the success of spore killing. Increasing the distance of *rfk-1* from the right border of *Sk-2* causes it to be inactivated by a genome defense process called meiotic silencing by unpaired DNA (MSUD), adding to accumulating evidence that MSUD exists, at least in part, to protect genomes from meiotic drive.

## INTRODUCTION

In eukaryotic organisms, genetic loci are typically transmitted through sexual reproduction to the next generation in a Mendelian manner. However, some loci possess the ability to improve their own transmission rate through meiosis at the expense of a competing locus. These “selfish” loci are often referred to as meiotic drive elements (Zimmering *et al.* 1970). The genomic conflict caused by meiotic drive elements may impact processes ranging from gametogenesis to speciation (Lindholm *et al*. 2016). Meiotic drive elements are found across the eukaryote tree of life (Burt and Trivers 2008; Bravo Núñez *et al*. 2018) and classic examples include *SD* in fruit flies (Larracuente and Presgraves 2012), the *t-*complex in mice (Lyon 2003; Sugimoto 2014), and *Ab10* in *Zea mays* (Rhoades 1952; Kanizay *et al.* 2013). In the fungal kingdom, the known meiotic drive elements achieve biased transmission through spore killing (Raju 1994) and a handful of spore killer systems have been studied in detail. While the prion-based spore killing mechanism of *het-s* in *Podospora anserina* is the best characterized (Dalstra *et al.* 2003; Saupe 2011), the mechanisms by which other fungal meiotic drive elements kill spores are mostly unknown (*e.g.*, see Grognet et al. 2014; Hu et al. 2017; Nuckolls et al. 2017).

Two fungal meiotic drive elements have been identified in the fungus *Neurospora intermedia* (Turner and Perkins, 1979). This species is closely related to the genetic model *Neurospora* c*rassa* (Davis 2000), and the mating processes in both fungi are essentially identical. Mating begins with fertilization of an immature fruiting body called a protoperithecium by a mating partner of the opposite mating type. After fertilization, the protoperithecium develops into a mature fruiting body called a perithecium. The nuclei from each parent multiply within the developing perithecium, and a single nucleus from each parent is sequestered into a tube-like meiotic cell (Raju 1980). Meiosis begins with fusion of the parental nuclei and ends with production of four recombinant daughter nuclei. Each recombinant nucleus proceeds through a single round of mitosis, resulting in a total of eight nuclei in the meiotic cell. A process known as ascosporogenesis then constructs cell walls and membranes around each nucleus to produce sexual spores called ascospores. Maturing ascospores accumulate a dark pigment and develop the shape of a spindle; thus, at the end of ascosporogenesis, the mature meiotic cells appear to contain eight miniature black American footballs (Figure 1A). The meiotic cells also serve as ascospore sacs (asci) and a single perithecium can produce hundreds of asci, each derived from a unique meiotic event.

**Figure 1.**
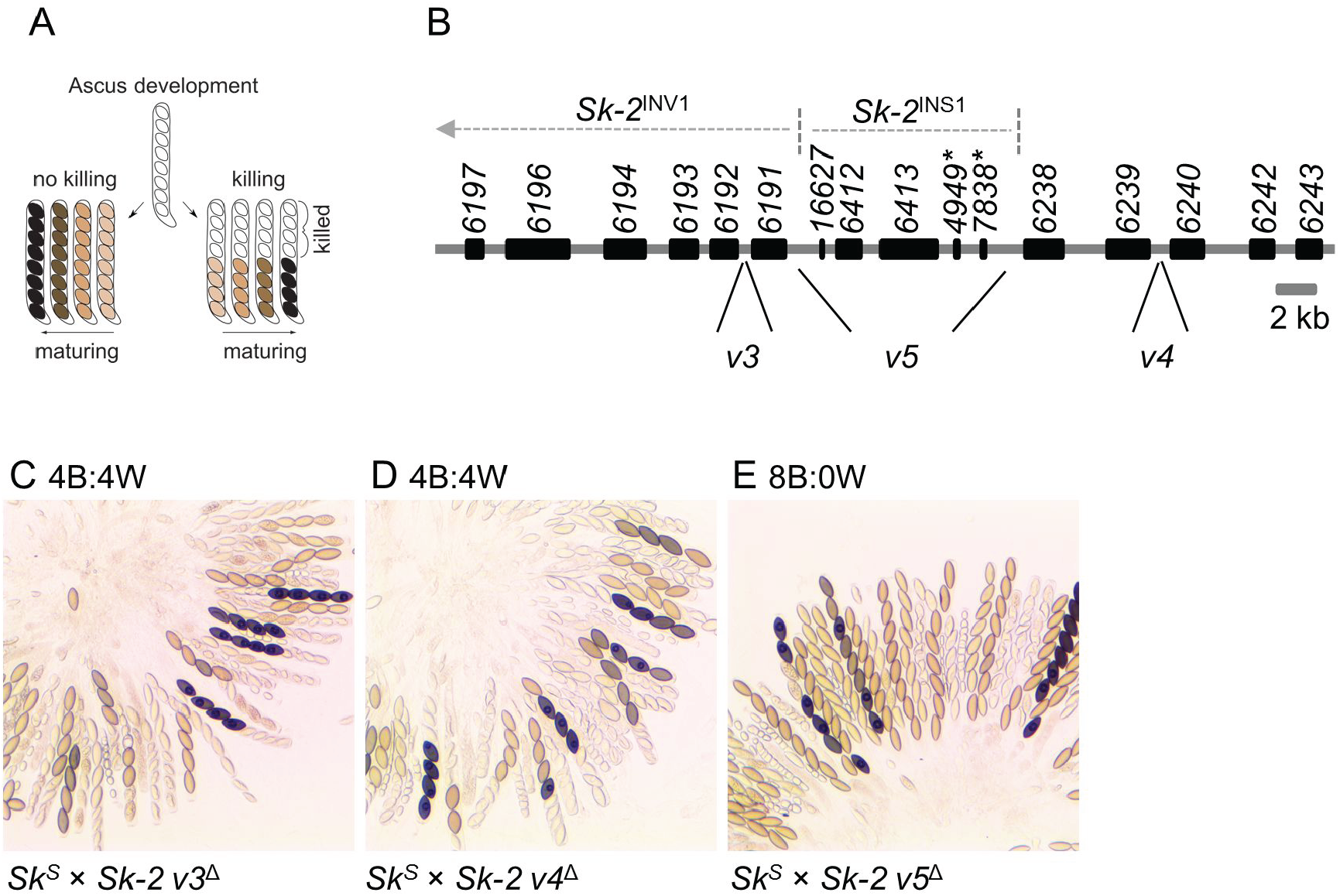
*Sk-2*^INS1^ harbors a genetic element required for spore killing. (A) The diagram illustrates phenotypic differences between spore killing and normal ascus development. Asci that have undergone spore killing contain four black and four white ascospores. Viable ascospores may appear tan, brown, or black, depending on their level of maturity. (B) Annotation of the *rfk*-*1* region as described by Harvey *et al.* (2014). Genes and pseudogenes are depicted as black rectangles. Gene names (*e.g., 6197*) are listed above the rectangles. Pseudogene names are appended with an asterisk. Labels *v3, v4*, and *v5* mark DNA intervals of the *rfk-1* region that were deleted and replaced with an *hph* selectable marker. (C–E) The images depict asci from crosses between *Sk*^*S*^ and *Sk-2* strains lacking different intervals of *Sk-2*^INS1^. The predominant phenotype of the asci produced by each cross is listed above each image. Crosses are as follows: (C) F2-23 × ISU-3023, (D) F2-23 × ISU-3017, and (E) F2-23 × ISU-3029.

During an effort in the 1970s to collect and characterize *Neurospora* isolates from around the world, Turner and Perkins discovered pairs of compatible mating partners that did not produce asci with eight viable ascospores (Perkins 1974; Turner and Perkins 1979). This outcome was more common when crosses were performed between isolates from widely separated populations, and in some cases the abnormal asci were attributed to heterozygosity of chromosome rearrangements between mating partners. However, for a few isolates of *N. intermedia*, asci with atypical phenotypes were due to chromosomal factors called *Spore killer-2* (*Sk-2*) and *Spore killer-3* (*Sk-3*). *Sk-2* and *Sk-3* are not single genes; rather, they are complexes of genes that span approximately 30 cM of chromosome III, and they are transmitted through meiosis as single units due to a recombination suppression mechanism thought to be enforced by inversions (Turner and Perkins 1979; Campbell and Turner 1987; Hammond *et al.* 2012; Harvey *et al.* 2014). Unlike standard genetic elements, which display a Mendelian transmission rate of 50% through sexual reproduction, *Sk-2* and *Sk-3* are transmitted at levels approaching 100% (Turner and Perkins 1979). This biased transmission occurs because *Sk-2* and *Sk-3* kill ascospores that do not inherit resistance to spore killing (Raju 1979; Turner and Perkins 1979). For example, in *Sk-2* × *Spore killer*-sensitive (*Sk*^*S*^) crosses, asci with four black ascospores and four clear (“white”) ascospores are produced (Figure 1A). This phenotype can be symbolized as 4B:4W. The four black ascospores are typically viable and nearly always of the *Sk-2* genotype, while the four white ascospores are inviable and presumed to be of the *Sk*^*S*^ genotype. The same phenomenon occurs in *Sk-3* × *Sk*^*S*^ crosses, except the four black ascospores are of the *Sk-3* genotype.

Although spore killers have not yet been detected in wild isolates of *N. crassa, Sk-2* and *Sk-3* have been introgressed into this species for genetic analysis. Introgression of *Sk-2* and *Sk-3* has allowed the discovery of resistance to spore killing in natural populations of *N. crassa* (Turner and Perkins 1979; Turner 2001). One of the *Sk-2*-resistant isolates (FGSC 2222) carries a resistant version of a gene whose function is best described by its name: *resistant to Spore killer* (*rsk*). Crosses of *rsk*^*LA*^ × *Sk-2*, where *rsk*^*LA*^ is the Louisiana allele of *rsk* carried by FGSC 2222, produce asci with an 8B:0W phenotype because ascospores inherit either *rsk*^*LA*^ or *Sk-2,* and both are sufficient for resistance to *Sk-2*-based spore killing (Hammond *et al.* 2012). Discovery of *rsk*^*LA*^ made identifying other *rsk* alleles possible, some of which do not provide resistance to the known spore killers. For example, the Oak Ridge *rsk* allele (*rsk*^*OR*^), typical of most laboratory strains, is resistant to neither *Sk-2* nor *Sk-3*. Additionally, some *rsk* alleles confer resistance to *Sk-3* but not *Sk-2*. An example is *rsk*^*PF5123*^, which exists in an *N. intermedia* isolate from French Polynesia. *Sk-2* and *Sk-3* also carry resistant versions of *rsk*, referred to as *rsk*^*Sk-2*^ and *rsk*^*Sk-3*^, respectively. Crosses homozygous for *Sk-2* (*i.e., Sk-2* × *Sk-2*) or *Sk-3* produce asci with an 8B:0W phenotype because each ascospore inherits a resistant *rsk* allele. Furthermore, heterozygous crosses between different spore killers (*e.g., Sk-2* × *Sk-3*) produce asci with a 0B:8W phenotype (Turner and Perkins 1979) because each ascospore inherits either *rsk*^*Sk-2*^ or *rsk*^*Sk-3*^ but not both (and *rsk*^*Sk-2*^ ascospores are killed by *Sk-3* while *rsk*^*Sk-3*^ ascospores are killed by *Sk-2*).

The Killer-Neutralization (KN) model has been proposed to explain how *Sk-2* and *Sk-3* achieve biased transmission through sexual reproduction (Hammond *et al.* 2012). The KN model holds that *Sk-2* and *Sk-3* each use a resistance protein and a killer protein (or nucleic acid) and both proteins are active throughout meiosis and ascosporogenesis. During the early stages of meiosis, in an *Sk*^*S*^ × *Sk-2* (or *Sk-3*) cross, both the resistance protein and the killer protein are hypothesized to diffuse throughout the meiotic cell. This unrestricted movement allows the resistance protein to neutralize the killer protein wherever the latter protein may be found. However, once ascospores are separated from the cytoplasm, the resistance protein becomes restricted to those ascospores that produce it (*e.g., Sk-2* ascospores), and ascospores that do not carry a resistant version of *rsk* (*e.g., Sk*^*S*^ ascospores) are subsequently killed. This model requires the killer protein to move between ascospores after ascospore delimitation or to have a long half-life that allows it to remain functional in sensitive ascospores.

Evidence for the KN model is seen in the outcome of *Sk*^*S*^ × *Sk-2 rsk*^Δ*Sk-2*^ crosses, where the latter strain has been deleted of its *rsk* allele. These crosses do not produce ascospores; instead, they produce asci that abort meiosis before ascospore production (Hammond *et al.* 2012). Meiotic cells of these crosses lack a resistant RSK, which likely causes the killing process to begin early in meiosis (at the ascus level) rather than during ascosporogenesis (at the ascospore level). The KN model is also supported by the existence of different *rsk* alleles. Previous studies have demonstrated the sequence of RSK to be the most important factor towards determining which killer it neutralizes (Hammond *et al.* 2012), suggesting that RSK and the killer may interact by a “lock and key” mechanism. To test this hypothesis, the killer must first be identified.

As described above, *Sk*^*S*^ × *Sk-2 rsk*^Δ*Sk-2*^ crosses produce abortive asci. We recently used this characteristic to screen for mutations that disrupt spore killing (Harvey *et al.* 2014). Specifically, we fertilized an *Sk*^*S*^ mating partner with mutagenized *Sk-2 rsk*^Δ*Sk-2*^ conidia (asexual spores that also function as fertilizing propagules). We reasoned that only an *Sk-2 rsk*^Δ*Sk-2*^ conidium mutated in a gene “*r*equired *f*or spore *k*illing” (*rfk*) would produce viable ascospores when crossed with *Sk*^*S*^. The screen allowed us to isolate six *rfk* mutants (ISU-3211 through ISU-3216). Complementation analysis of each mutant strain suggested all to be mutated at the same locus, which was subsequently named *rfk-1* and mapped to a 45 kb region within *Sk-2* on chromosome III. Here, we report the identification of *rfk-1* as a gene encoding a protein of at least 39 amino acids. In addition to identifying *rfk-1*, we have found that the cellular process of meiotic silencing by unpaired DNA places limits on the location of *rfk-1* within *Sk-2*. The implications of this finding with respect to meiotic drive element evolution are discussed.

## MATERIALS AND METHODS

### Strains, media, and crossing conditions

The strains used in this study are listed along with genotype information in Table 1. Vogel’s minimum medium (Vogel 1956), with supplements as required, was used to grow and maintain all strains. Hygromycin B and nourseothricin sulfate (Gold Biotechnology) were used at a working concentration of 200 μg / ml and 45 μg / ml, respectively. Synthetic crossing medium (pH 6.5) with 1.5% sucrose, as described by Westergaard and Mitchell (1947), was used for crosses. Crosses were unidirectional and performed on a laboratory benchtop at room temperature under ambient lighting (Samarajeewa *et al.* 2014). After fertilization, crosses were allowed to mature for 12-16 days before perithecial dissection in 25 or 50% glycerol and asci were examined with a standard compound light microscope and imaging system. Ascus phenotype designations were based on qualitative observations. More than 90% of the asci from a cross had to display the same phenotype to receive one of the following designations: 8B:0W, 4B:4W, or aborted.

**Table 1.**
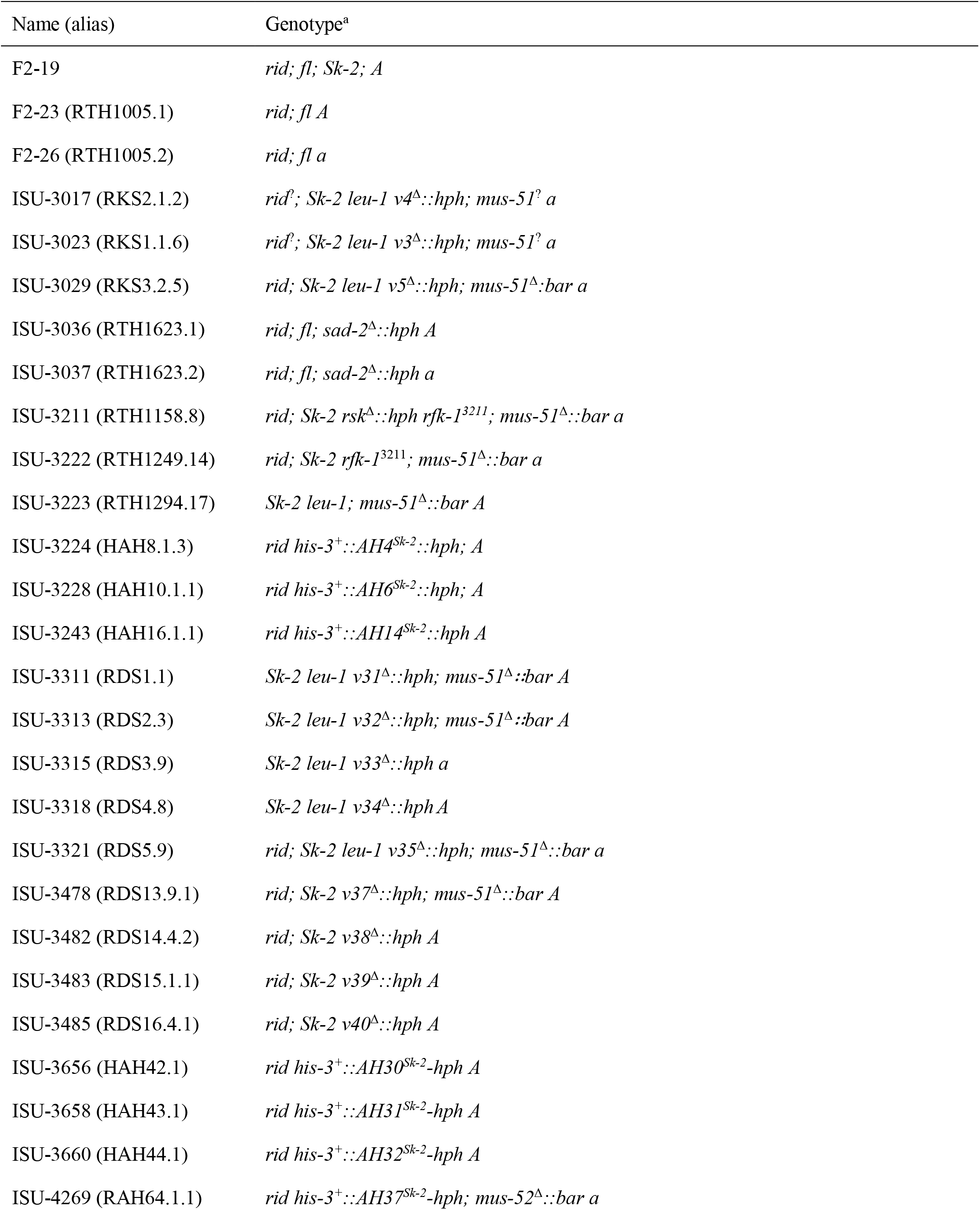
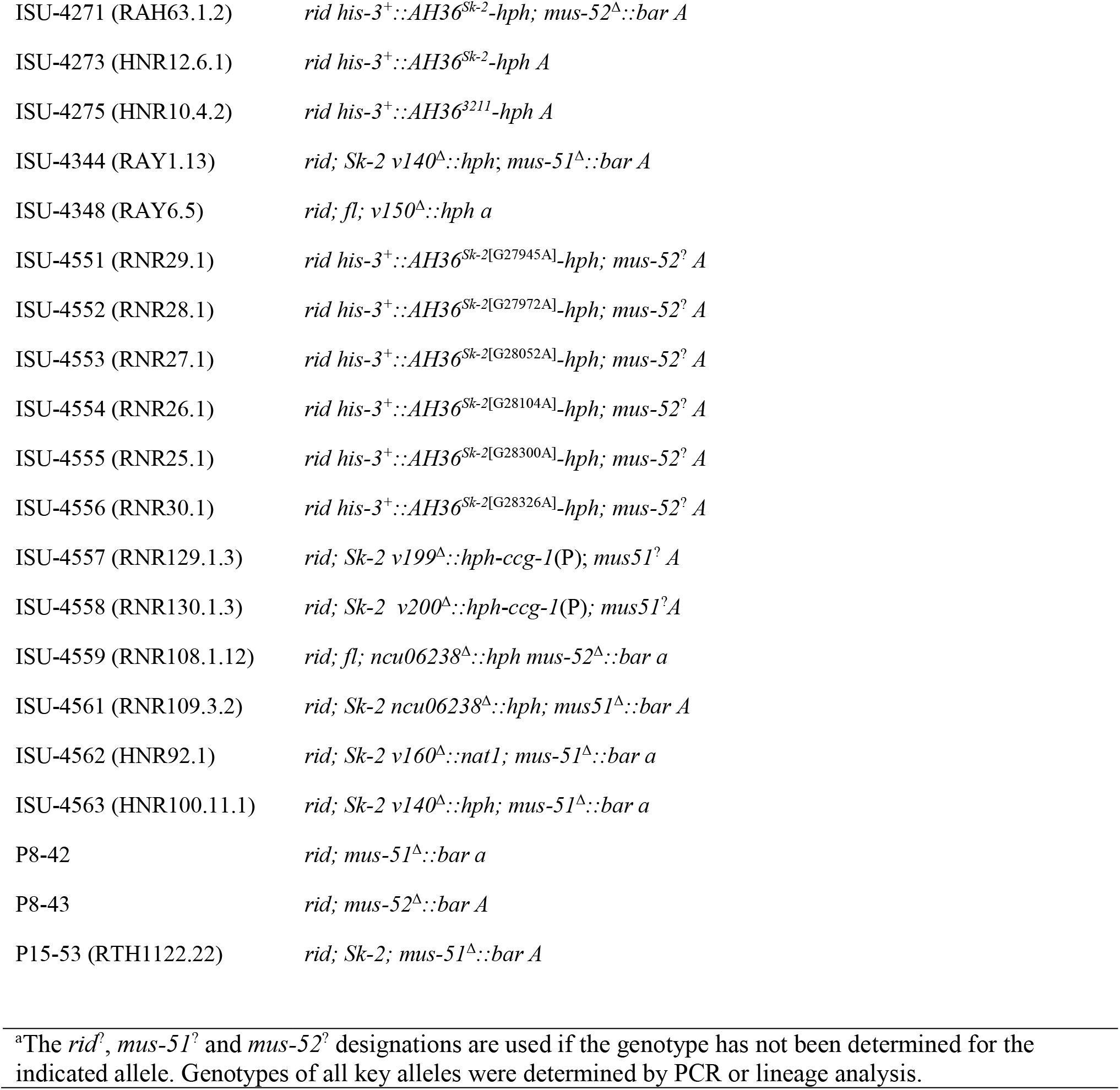
Strains used in this study

### Genetic modification of *N. crassa*, genotyping, and sequence confirmations

A technique called double-joint PCR was used to construct all deletion vectors (Yu *et al.* 2004; Hammond *et al.* 2011). Transgene-insertion vectors were designed to insert transgenes along with a hygromycin resistance cassette (*hph*) next to *his-3* on chromosome I. Construction details for deletion and insertion vectors are provided in Supporting Information (Tables S1–S4). Transformations of *N. crassa* were performed by electroporation of conidia (Margolin *et al*. 1997). Homokaryons were derived from heterokaryotic transformants with a microconidium isolation technique (Ebbole and Sachs, 1990) or by crossing the transformants to standard laboratory strains (F2-23 or F2-26) to obtain homokaryotic ascospores. Site-directed mutagenesis was performed essentially as described for the QuikChange II Site-Directed Mutagenesis Kit (Revision E.01, Agilent Technologies) and details for its use are provided in Table S5. All genotypes were confirmed by polymerase chain reaction (PCR) assays on genomic DNA isolated from lyophilized (freeze-dried) mycelia with IBI Scientific’s Mini Genomic DNA Kit (Plant/Fungi). Sanger sequencing was used to confirm sequences and/or identify mutations in PCR products and plasmids.

### Data availability

All strains and plasmids generated during this study are available upon request. Supplemental files available at FigShare.

## RESULTS

### Deletion of a DNA interval spanning most of *Sk-2*^INS1^ eliminates spore killing

The annotated 45 kb *rfk-1* region contains 14 protein-coding genes, two pseudogenes (denoted with an asterisk), an inverted sequence (*Sk-2*^INV1^), an inversion breakpoint, and an 11 kb insertion sequence (*Sk-2*^INS1^; GenBank: KJ908288.1; Figure 1B). To refine the location of *rfk-1* within this 45 kb region, intervals *v3, v4*, and *v5* (Figure 1B and Table 2) were deleted and replaced with *hph* and the resulting deletion strains were crossed with an *Sk*^*S*^ mating partner. We found that while deletion of interval *v3* or *v4* had no effect on spore killing (asci are 4B:4W; Figure 1, C and D), deletion of *v5* eliminated it (asci are 8B:0W; Figure 1E).

**Table 2.**
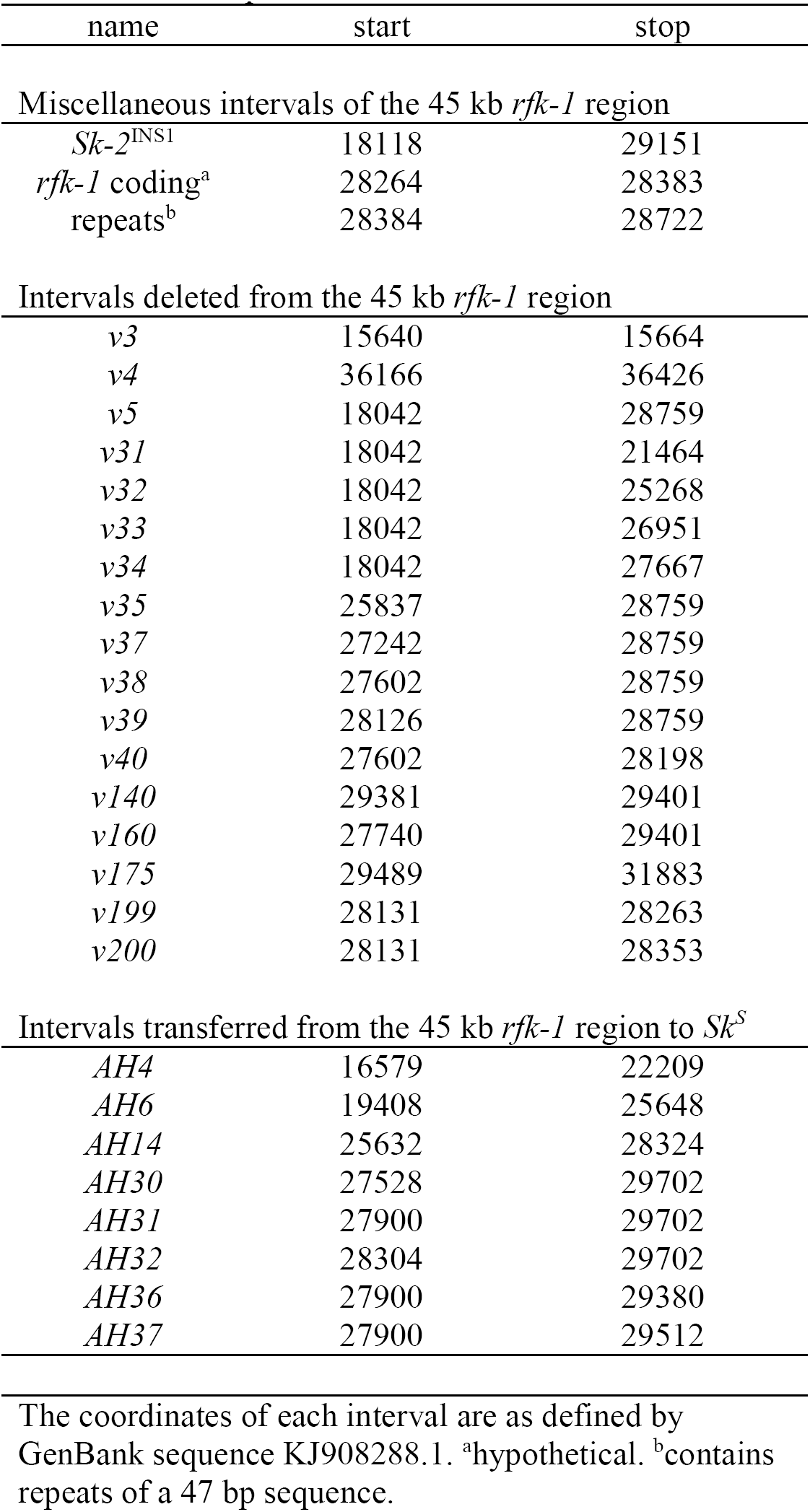
Interval positions

### A DNA interval between *ncu07838*^*\**^ and *ncu06238* is required for spore killing

Interval *v5* spans most of *Sk-2*^INS1^ (Figure 2, A and B). To further refine the position of *rfk-1* within *Sk-2*^INS1^, we constructed nine additional deletion strains and crossed each one with an *Sk*^*S*^ mating partner (Figure 2B and Table 2). Surprisingly, deletion of the annotated genes and pseudogenes within *Sk-2*^INS1^ did not interfere with spore killing (Figure 3, A–D). In contrast, deletion of the intergenic region between *ncu07838\** and *ncu06238* eliminated spore killing (Figure 3, E–I).

**Figure 2.**
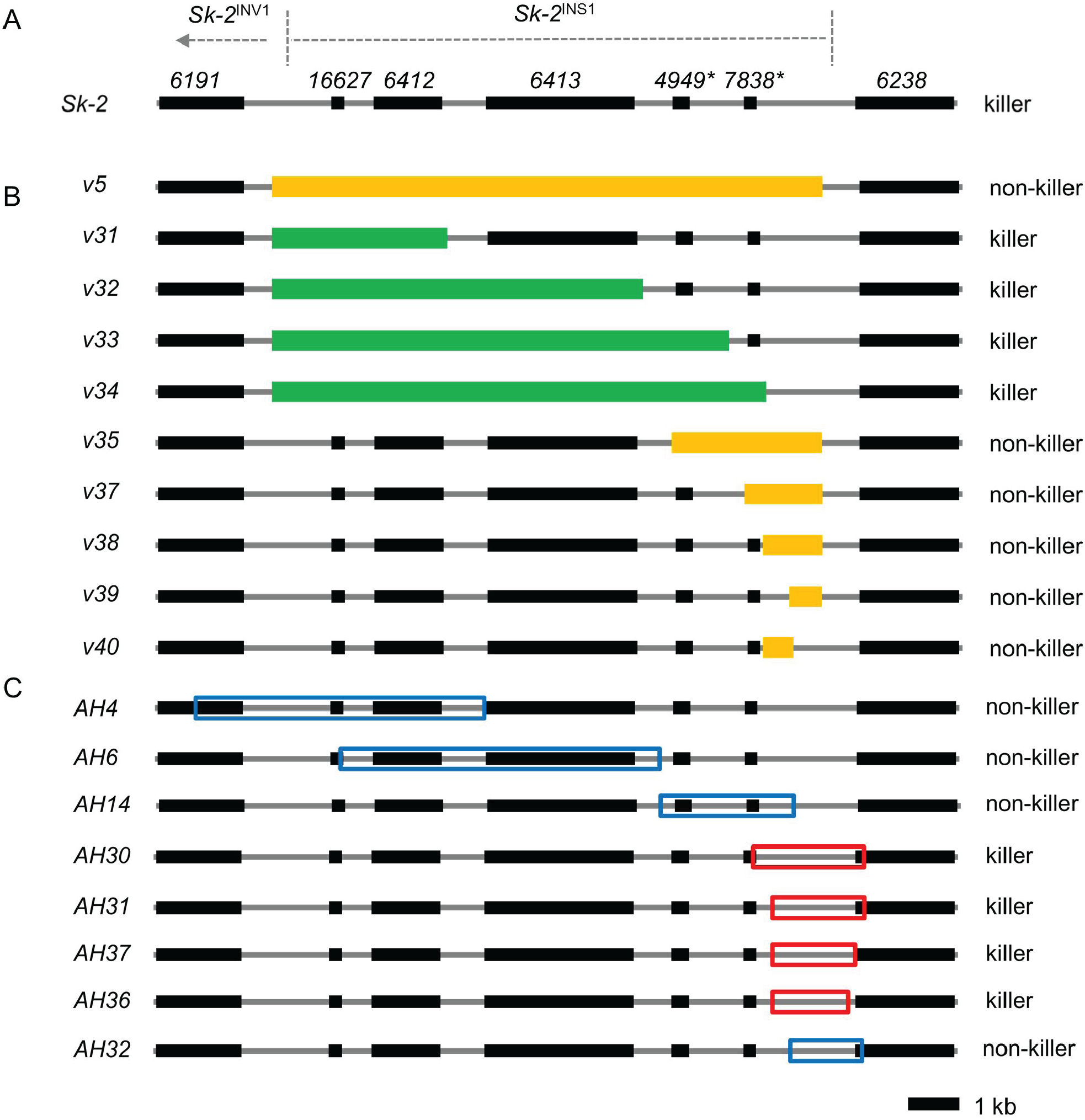
Deletion and insertion maps. (A) A diagram of *Sk-2*^INS1^. (B) Ten intervals of *Sk-2*^INS1^ were deleted and replaced with *hph*. For convenience, each interval is named according to its deletion vector (*e.g.*, interval *v5* is named after deletion vector v5). Orange rectangles mark intervals that disrupt spore killing upon deletion. Green rectangles mark intervals that do not disrupt spore killing when deleted. (C) Eight intervals of *Sk-2*^INS-1^were transferred to the *his-3* locus of an *Sk*^*S*^ strain. Intervals were named according to the name of the plasmid used for cloning of the interval (*e.g.*, interval *AH4* is named after plasmid pAH4). Red and blue open rectangles mark killer (*i.e.*, abortion-inducing) and non-killer intervals, respectively.

**Figure 3.**
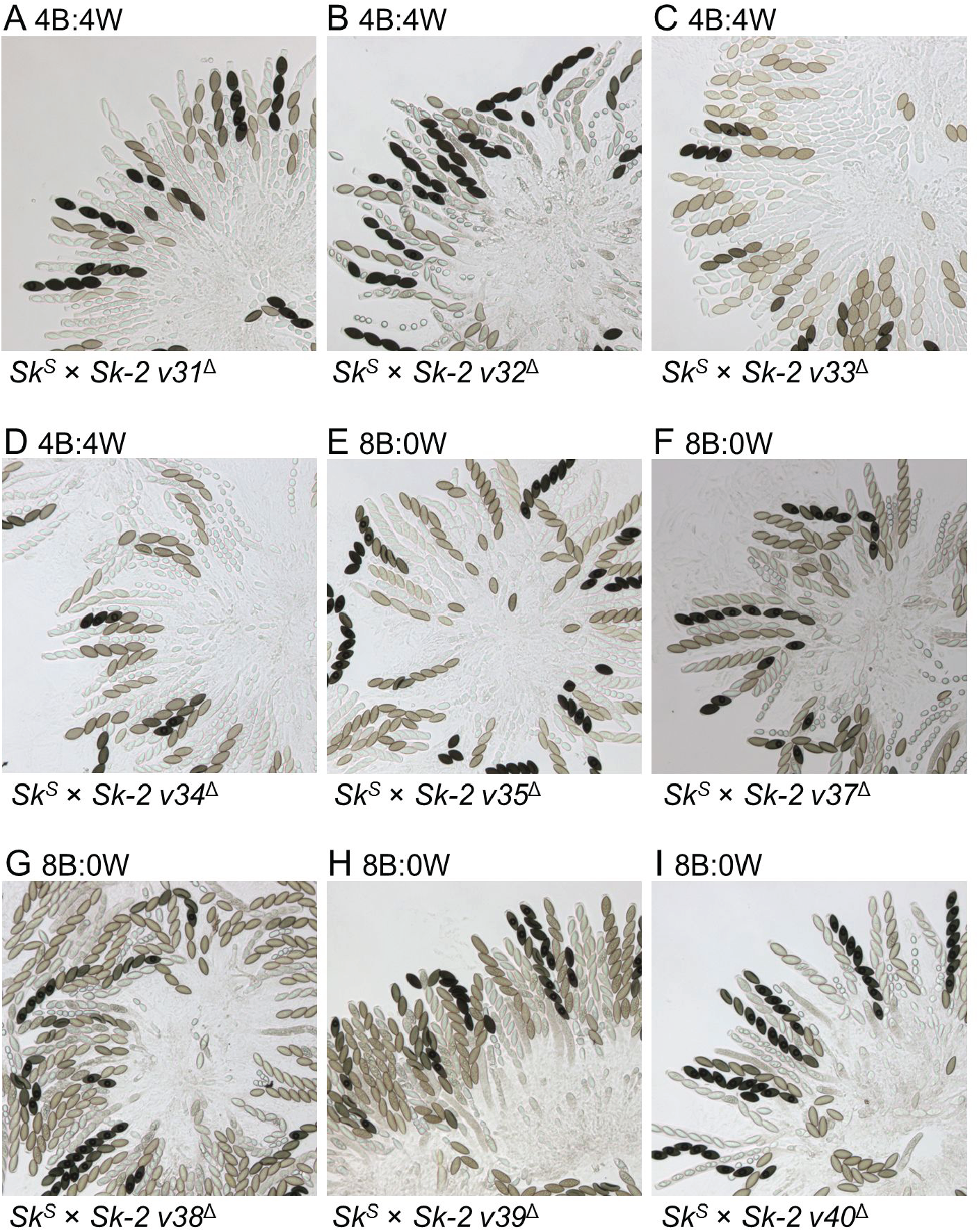
Deletion of a genetic element between pseudogene *7838*^*^ and the right border of *Sk-2*^INS1^ eliminates spore killing. (A–I) The images depict asci from crosses between an *Sk*^*S*^ strain and an *Sk-2* mating partner lacking a specific interval of *Sk-2*^INS1^. Crosses are as follows: (A) F2-26 × ISU-3311, (B) F2-26 × ISU-3313, (C) F2-23 × ISU-3315, (D) F2-26 × ISU-3318, (E) F2-23 × ISU-3321, (F) F2-26 × ISU-3478, (G) F2-26 × ISU-3482, (H) F2-26 × ISU-3483, and (I) F2-26 × ISU-3485.

### An ascus aborting element exists between *ncu07838*^*\**^ and *ncu06238*

The above results suggest that *rfk-1* is found within the intergenic region between *ncu07838*^*^ and *ncu06238* and that *rfk-1* is required for spore killing. But, is *rfk-1* also sufficient for spore killing? To answer this question, we genetically-modified eight *Sk*^*S*^ strains to carry different intervals of *Sk-2*^INS1^ (Figure 2C and Table 2) and found that each strain produced normal asci when crossed with an *Sk*^*S*^ mating partner (Figure S1). The reason for this finding can be traced to a silencing process called meiotic silencing by unpaired DNA (MSUD; Hammond 2017; Aramayo and Selker 2013). In a standard cross, where only one mating partner carries an ectopic transgene (*e.g.*, an interval of *Sk-2*^INS1^), MSUD identifies the transgene as unpaired and silences it for the duration of meiosis. Therefore, to detect a phenotype that requires the expression of an unpaired transgene during meiosis, it is often necessary to suppress MSUD. MSUD suppression can be achieved by deleting a gene called *sad-2* from one mating partner of a cross (Shiu *et al.* 2006). With this technique, we found that some *Sk-2*^INS1^ intervals have no effect on ascus development, while others abort it. For example, normal asci are produced by strains carrying intervals *AH4*^*Sk-2*^, *AH6*^*Sk-2*^, *AH14*^*Sk-2*^, or *AH32*^*Sk-2*^ (Figure 4, A–C and H), while aborted asci are produced by strains carrying intervals *AH30*^*Sk-2*^, *AH31*^*Sk-2*^, *AH36*^*Sk-2*^, or *AH37*^*Sk-2*^ (Figure 4, D–G). The ascus abortion phenotype can be explained by the presence of *rfk-1* without the presence of a resistant version of *rsk*. Taken together, these findings suggest that intervals *AH30*^*Sk-2*^, *AH31*^*Sk-2*^, *AH36*^*Sk-2*^, and *AH37*^*Sk-2*^ contain *rfk-1* and that *rfk-1* is sufficient for spore killing.

**Figure 4.**
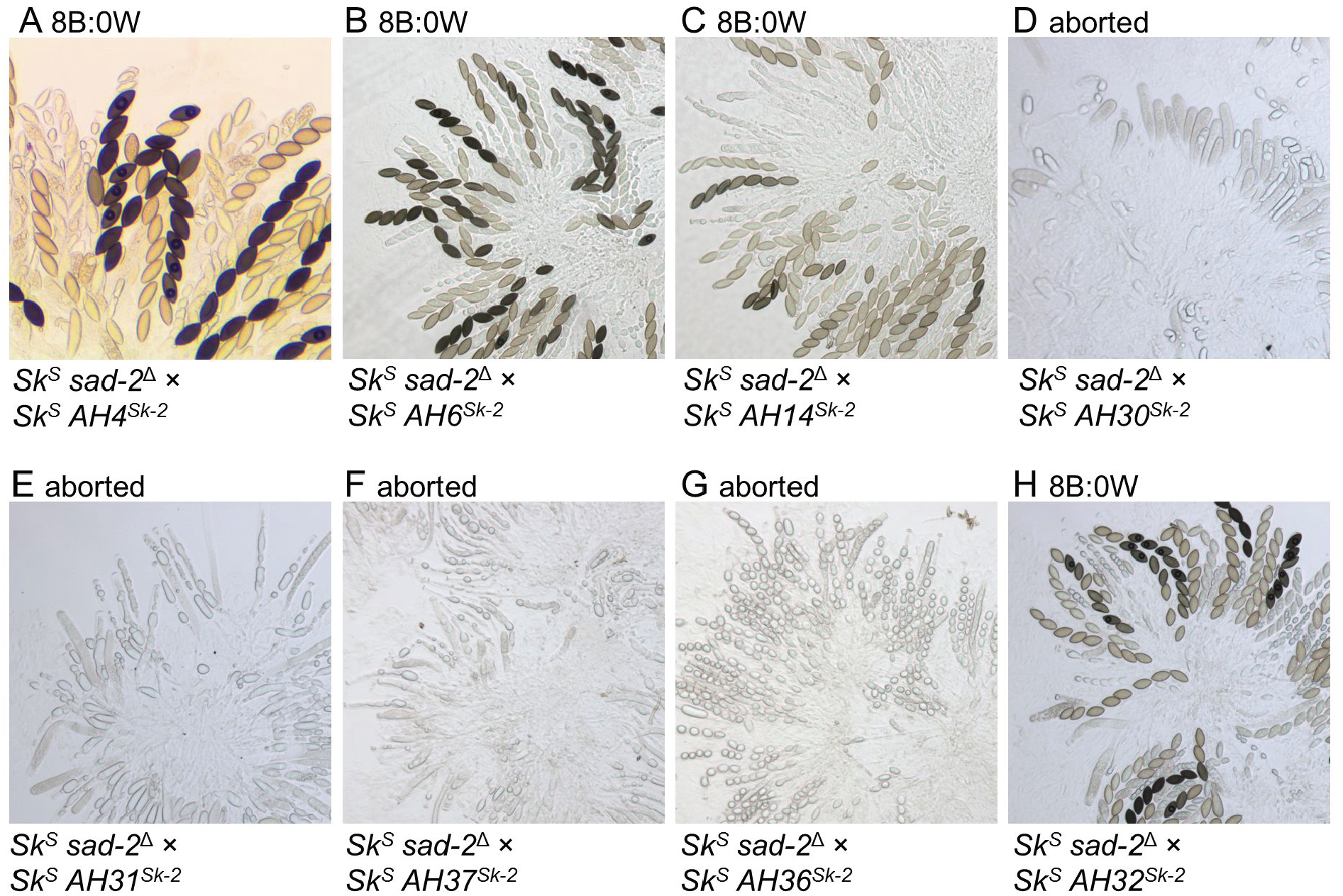
A genetic element within *Sk-2*^INS1^ causes ascus abortion upon its transfer to an *Sk*^*S*^ strain. (A–H) Images depict asci from crosses between an *Sk*^*S*^ *sad-2*^Δ^ strain and an *Sk*^*S*^ mating partner carrying a specific interval of *Sk-2*^INS1^. Crosses are as follows: (A) ISU-3037 × ISU-3224, (B) ISU-3037 × ISU-3228, (C) ISU-3036 × ISU-3243, (D) ISU-3037 × ISU-3656, (E) ISU-3037 × ISU-3658, (F) ISU-3036 × ISU-4269, (G) ISU-3037 × ISU-4271, and (H) ISU-3037 × ISU-3660.

### The *AH36* interval from an *rfk-1* strain does not cause ascus abortion

The shortest abortion-inducing interval identified by the above experiments is *AH36*, located between positions 27,899 and 29,381 of the 45 kb *rfk-1* region (Figure 2C and Table 2). Because the research path that led us to *AH36* began with mapping the position of *rfk-1* in strain ISU-3211 (Harvey *et al.* 2014), *AH36* in ISU-3211 (referred to as *AH36*^3211^) should harbor at least one mutation that disrupts *rfk-1* function. To test this hypothesis, we transferred *AH36*^3211^ to an *Sk*^*S*^ genetic background and crossed the resulting strain with an *Sk*^*S*^ *sad-2*^Δ^ mating partner. As expected, we found that *Sk*^*S*^ *sad-2*^Δ^ × *Sk*^*S*^ *AH36*^3211^ crosses produce normal asci (Figure 5).

**Figure 5.**
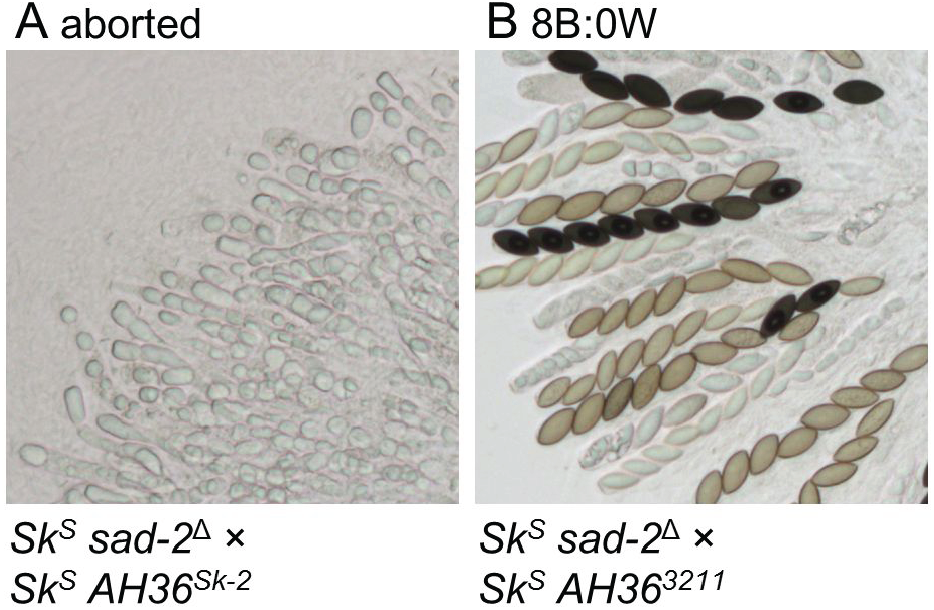
The *AH36* interval from an *rfk-1* mutant does not cause ascus abortion. Images depict asci from crosses between an *Sk*^*S*^ *sad-2*^Δ^ strain and an *Sk*^*S*^ strain carrying either the *AH36* interval from (A) F2-19 (*rfk-1*^*+*^) or (B) ISU-3211 (an *rfk-1* mutant). Crosses are as follows: (A) ISU-3037 × ISU-4273 and (B) ISU-3037 × ISU-4275.

### The G28326A mutation disrupts the ascus-aborting ability of *AH36*^*Sk-2*^

The different phenotypes associated with *AH36*^*Sk-2*^ and *AH36*^3211^ suggest that they differ at the sequence level. Indeed, sequencing of these two alleles allowed us to identify seven guanine to adenine transition mutations in *AH36*^3211^ (Figure 6A; G27904A, G27945A, G27972A, G28052A, G28104A, G28300A, and G28326A). To determine if one (or more) of these mutations is responsible for the inability of *AH36*^3211^ to cause ascus abortion, we examined six of the seven mutations by site-directed mutagenesis. For each mutation, this involved mutating the base in a clone of interval *AH36*^*Sk-2*^, placing the mutated interval (*e.g., AH36*^*Sk-^2^* [G27945A]^) in an *Sk*^*S*^ strain, and crossing the transgenic strain to an *Sk*^*S*^ *sad-2*^Δ^ mating partner. Through this procedure, we found that only one of the six mutations examined (*i.e.,* G28326A) eliminates the ascus-aborting ability of *AH36*^*Sk-2*^ (Figure 7).

**Figure 6.**
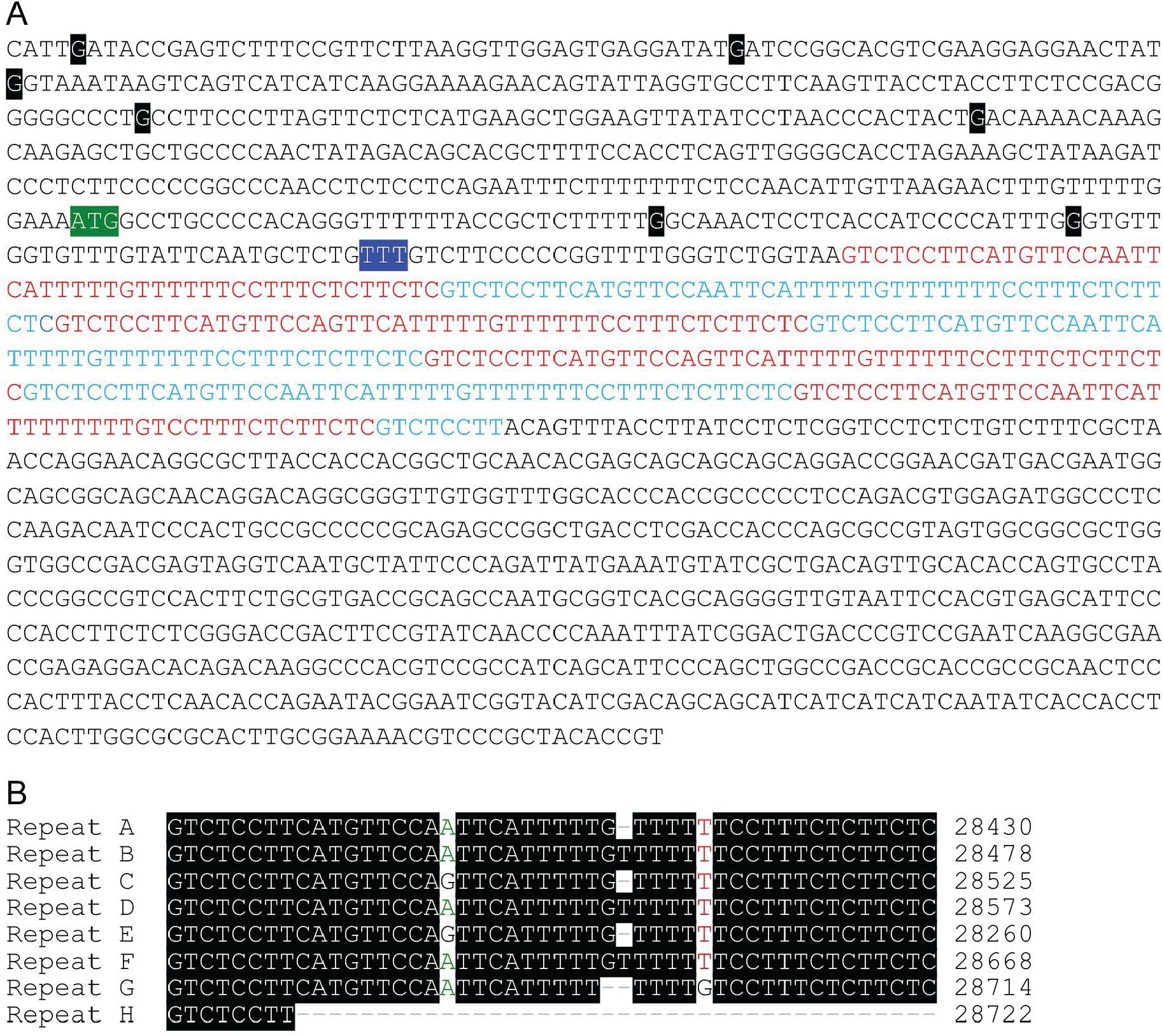
The *AH36* interval from an *rfk-1* mutant contains seven point mutations. (A) The 1481 bp sequence of *AH36*^*Sk-2*^ is shown. A region containing 7.17 repeats of a 46–48 bp sequence is highlighted with red and blue fonts. The colors alternate with each iteration of the repeated sequence. The *AH36*^3211^ sequence contains seven G to A transition mutations. The position of each mutation is marked by a white character on a black background with the non-mutated character shown. The mutations in *AH36*^3211^ are (from left to right): G27904A, G27945A, G27972A, G28052A, G28104A, G28300A, and G28326A. A non-native promoter was placed directly upstream of a putative RFK-1 start codon (white font, green background) with vector v199. A non-native promoter was also placed directly upstream of a TTT triplet (white font, blue background) with vector v200. (B) Alignment of the repetitive sequences highlighted in panel A.

**Figure 7.**
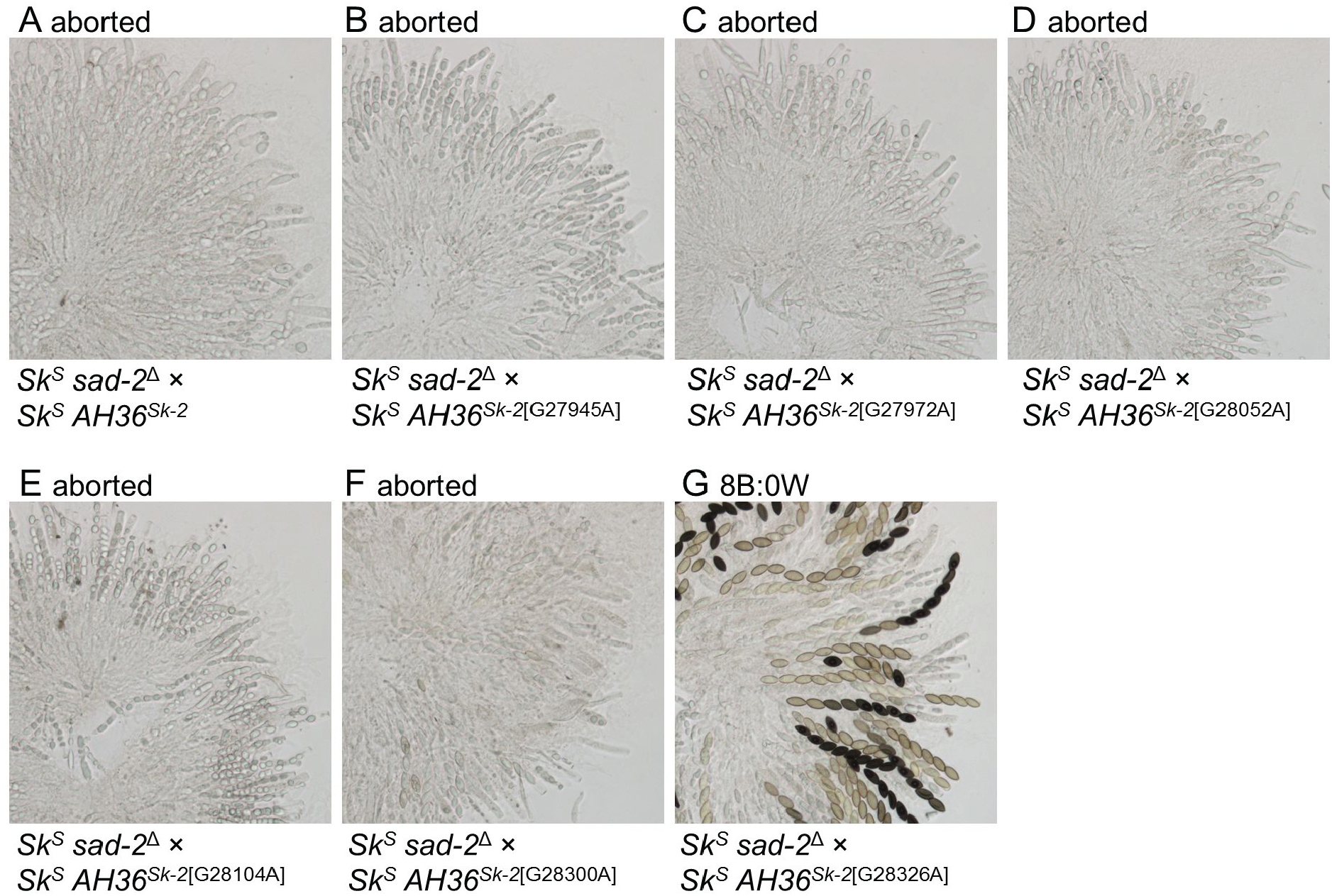
A point mutation within *AH36* eliminates its ability to abort ascus development. Six of the seven point mutations in *AH36*^3211^ were examined for a potential role in ascus abortion. Each mutation was placed individually in *AH36*^*Sk-2*^ by site-directed mutagenesis. (A–G) Images depict asci from crosses between an *Sk*^*S*^ *sad-2*^Δ^ strain and an *Sk*^*S*^ mating partner carrying *AH36*^*Sk-2*^ or one of its mutated derivatives. Crosses are as follows: (A) ISU-3037 × ISU-4273, (B) ISU-3037 × ISU-4551, (C) ISU-3037 × ISU-4552, (D) ISU-3037 × ISU-4553, (E) ISU-3037 × ISU-4554, (F) ISU-3037 × ISU-4555, (G) ISU-3037 × ISU-4556.

We also identified a 46–48 bp tandem repeat (7.17 repeats) between positions 28,384 and 28,722 (Figure 6, A and B). The sequences of *AH36*^*Sk-2*^ and *AH36*^3211^ are identical between these positions and thus the biological significance of the tandem repeats with respect to spore killing is currently unknown.

### A putative start codon for RFK-1 is located within *AH36*

The G28326A mutation is 62 bp to the right of a putative start codon at position 28,264 (Figure 6). To test if this “ATG” could serve as the start codon for RFK-1, we constructed two deletion vectors: v199 and v200 (Figure 8A). Vector v199 deletes the interval between 28,131 and 28,264 and replaces it with *hph* and the promoter of the *N. crassa ccg-1* gene, thereby inserting *hph-ccg-1*(P) directly upstream of the ATG at position 28,264 (Figure 8B). As a control, we used vector v200 to place *hph-ccg-1*(P) directly upstream of position 28,354, located 90 bases to the right of the proposed *rfk-1* start codon. When inserted directly upstream of 28,264, *hph-ccg-1*(P) has no effect on spore killing (Figure 8, C and D). In contrast, when inserted 90 bases to the right of this position, *hph-ccg-1*(P) disrupts spore killing (Figure 8E). These findings demonstrate that the ATG at position 28,264 could serve as the *rfk-1* start codon. Furthermore, they suggest that placement of *hph*-*ccg-1*(P) directly upstream of position 28,354 interrupts the *rfk-1* coding region.

**Figure 8.**
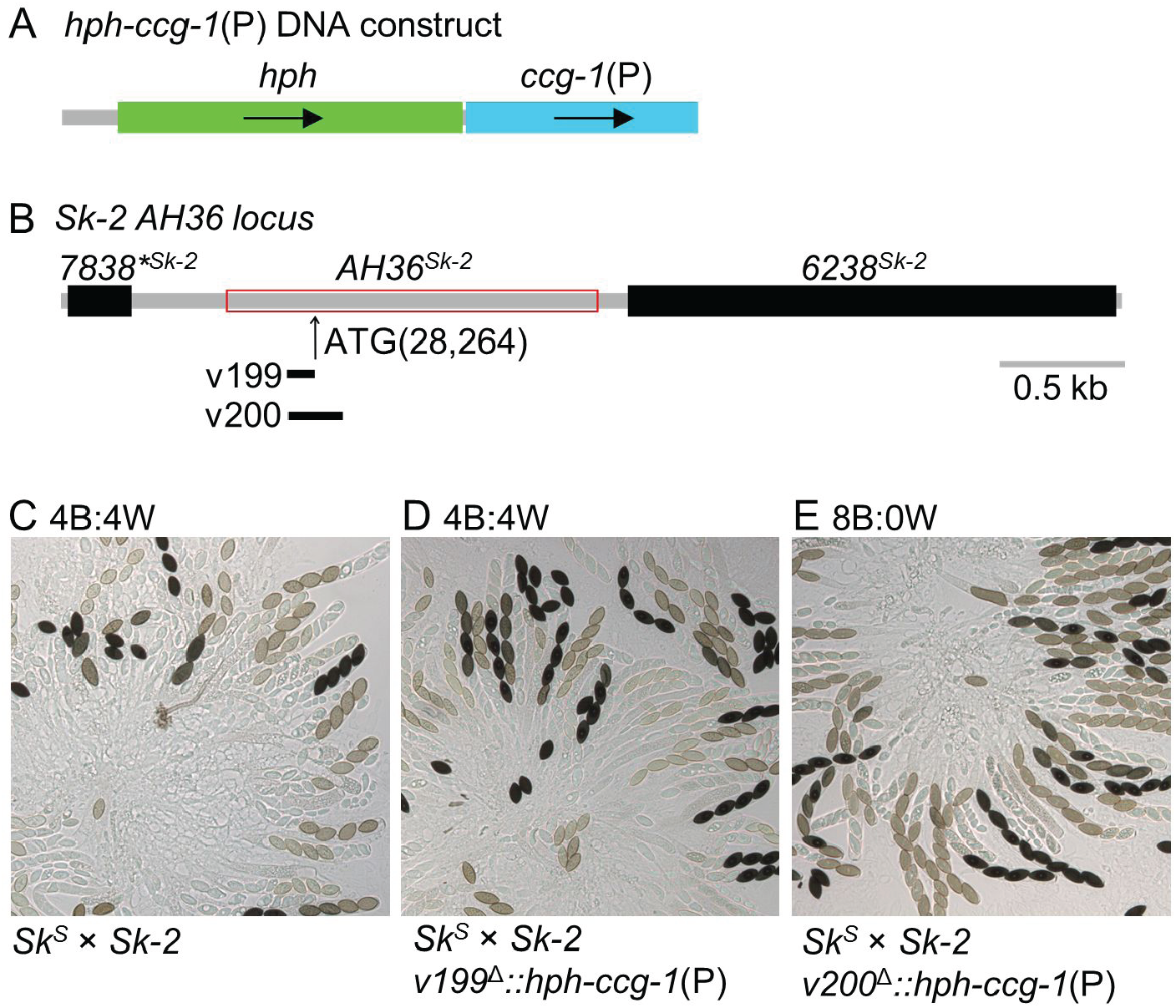
A putative start codon for RFK-1 exists within *AH36*. (A) A DNA construct consisting of *hph* and the promoter for the *N. crassa ccg-1* gene was used to make two transformation vectors. (B) Vector 199 (v199) was designed to replace 133 bp of *AH36* while fusing *ccg-1*(P) to a putative ATG codon at position 28,264 (Figure 6), thereby creating the *v199*^*Δ*^::*hph-ccg-1*(P) allele. Similarly, vector 200 (v200) was designed to replace 223 bp of *AH36* while fusing *ccg-1*(P) to position 28,354 (Figure 6), thereby creating the *v200*^*Δ*^::*hph-ccg-1*(P) allele. (C–E) Crosses were performed to determine the effect of each allele on spore killing. Images depict asci from the following crosses (C) F2-26 × P15-53, (D) F2-26 × ISU-4557, and (E) F2-26 × ISU-4558.

### The arrangement of *rfk-1* within *Sk-2* protects it from MSUD

The right border of *Sk-2* is found at position 29,151 (Figure 9A, dotted line; Table 2; Harvey *et al.* 2014). To the right of this position, the sequences of *Sk-2* and *Sk*^*S*^ strains are very similar. For example, a simple ClustalW alignment (Thompson *et al.* 1994; Hall 1999) finds that *Sk-2* positions 29,152 through 35,728 are 94.4% identical to the corresponding positions within *Sk*^*S*^ (GenBank: CM002238.1, positions 2,011,073 to 2,017,662). In contrast, the sequences to the left of the *Sk-2* border are unrelated between *Sk-2* and *Sk*^*S*^ strains (Figure 9A). Interestingly, most of *AH36* is found to the left of the *Sk-2* border, and thus most of *AH36*, including *rfk-1*, is unpaired during meiosis in *Sk*^*S*^ × *Sk-2* crosses. If so, how does *rfk-1* avoid inactivation by MSUD? While the molecular details of how MSUD detects unpaired DNA are unknown, we considered the possibility that the distance of *rfk-1* from a “paired” sequence allows it to avoid MSUD (*e.g.*, see the *ncu06238* genes in *Sk-2* and *Sk*^*S*^, Figure 9A). To test this hypothesis, we inserted *hph* immediately to the right of *AH36* in a standard *Sk-2* strain (Figure 9A). We refer to this particular allele as *v140*^Δ^::*hph*. The *v140*^Δ^::*hph* allele increases the distance of *rfk-1* from paired sequences by a length of 1391 bp (the length of *hph* minus the 21 bp that were deleted by v140). As predicted, we found that spore killing is absent in *Sk*^S^ × *Sk-2 v140*^Δ^::*hph* crosses (Figure 9B). To confirm that the lack of spore killing is a result of the increased distance of *rfk-1* from paired DNA during meiosis, we inserted *hph* at the corresponding location in an *Sk*^*S*^ strain (Figure 9A). We refer to this allele as *v150*^Δ^::*hph*. When an *Sk*^*S*^ *v150*^Δ^::*hph* strain is crossed with an *Sk-2 v140*^Δ^::*hph* strain, spore killing is normal (Figure 9C). Thus, the proximity of *rfk-1* to paired DNA helps it avoid inactivation by MSUD. As a final test of this hypothesis, we crossed *Sk*^S^ *sad-2*^Δ^ and *Sk-2 v140*^Δ^::*hph* mating partners and found that spore killing is also normal in this cross (Figure 9D), most likely because *sad-2*^Δ^ suppresses MSUD, which makes the distance of *rfk-1* from paired sequences irrelevant to the expression of *rfk-1* during meiosis.

**Figure 9.**
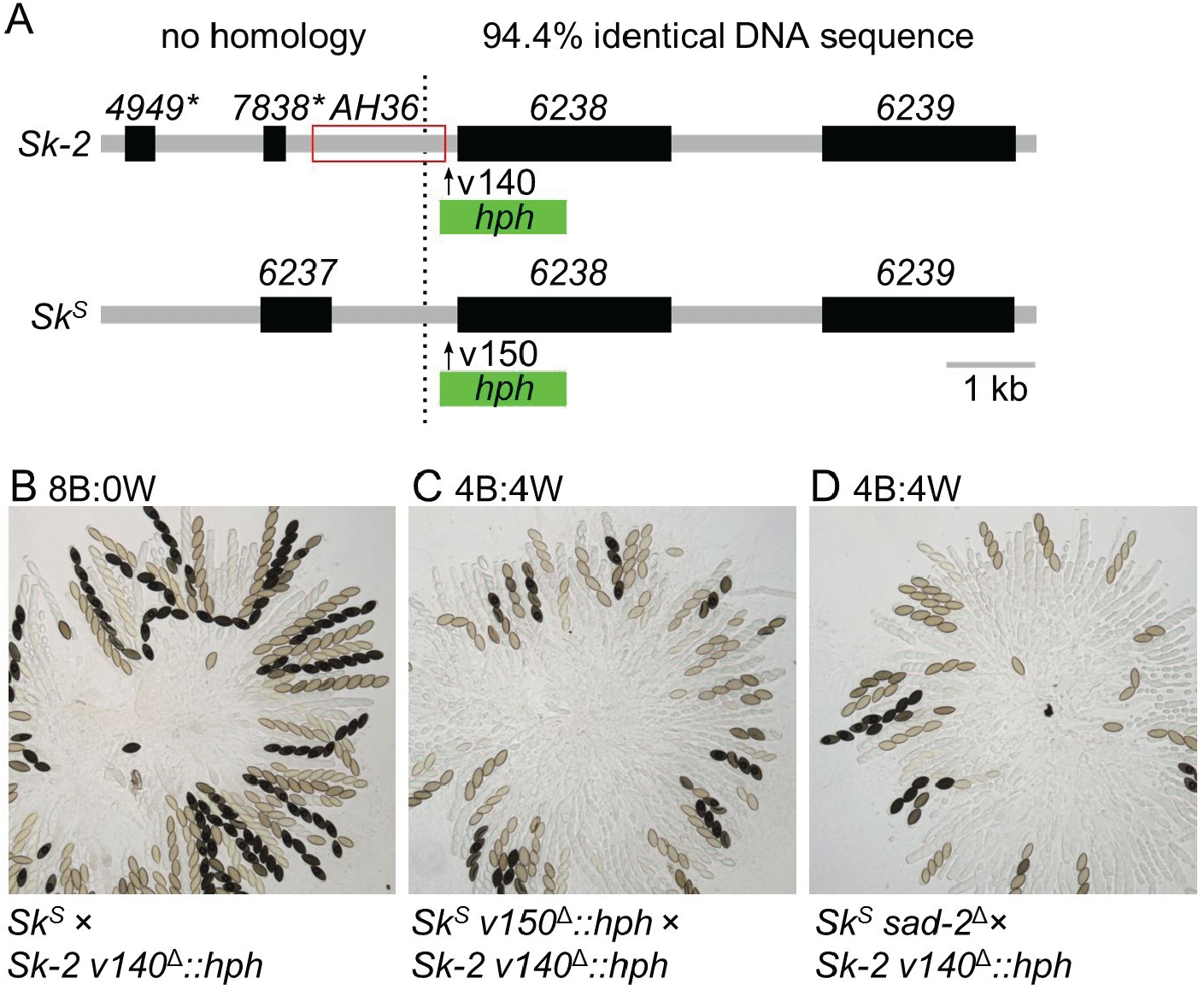
The native arrangement of *rfk-1* protects it from MSUD. (A) Interval *AH36* spans the right border of *Sk-2* (marked by a black dotted line). An *hph* selectable marker was placed immediately to the right of *AH36* in an *Sk-2* strain (with vector v140) to create the *v140*^Δ^::*hph* allele and at the corresponding location in an *Sk*^*S*^ strain (with vector v150) to create the *v150*^Δ^::*hph* allele. (B–D) Crosses were performed to determine the effect of each allele on spore killing. Images depict asci from the following crosses: (B) F2-23 × ISU-4344, (C) ISU-4348 × ISU-4344, and (D) ISU-3036 × ISU-4344.

### The *rfk-1* gene does not include *ncu06238*

To confirm that *ncu06238*, the gene to the right of *rfk-1* (as depicted in Figure 10A), is not required for spore killing, we deleted *ncu06238* from both *Sk*^*S*^ and *Sk-2* and analyzed ascus phenotypes in crosses involving *ncu06238* deletion strains. However, we found that *Sk*^*S*^ *ncu06238*^Δ^ × *Sk*^*S*^ crosses produce asci with varying numbers of fully developed ascospores (Figure 10A). Therefore, we could not use ascus phenotype to determine if spore killing is functional in *Sk*^*S*^ *ncu06238*^Δ^ × *Sk-2 ncu06238*^Δ^ crosses (Figure 10B). Instead, we calculated the percentage of progeny with an *Sk-2* genotype produced by a cross between *Sk*^*S*^ *ncu06238*^Δ^ and *Sk-2 ncu06238*^Δ^ mating partners. We found that 46 of 47 progeny had the *Sk-2* genotype (data not shown). Therefore, because meiotic drive functions without *ncu06238*, the *rfk-1* coding region does not overlap or include positions occupied by *ncu06238.*

**Figure 10.**
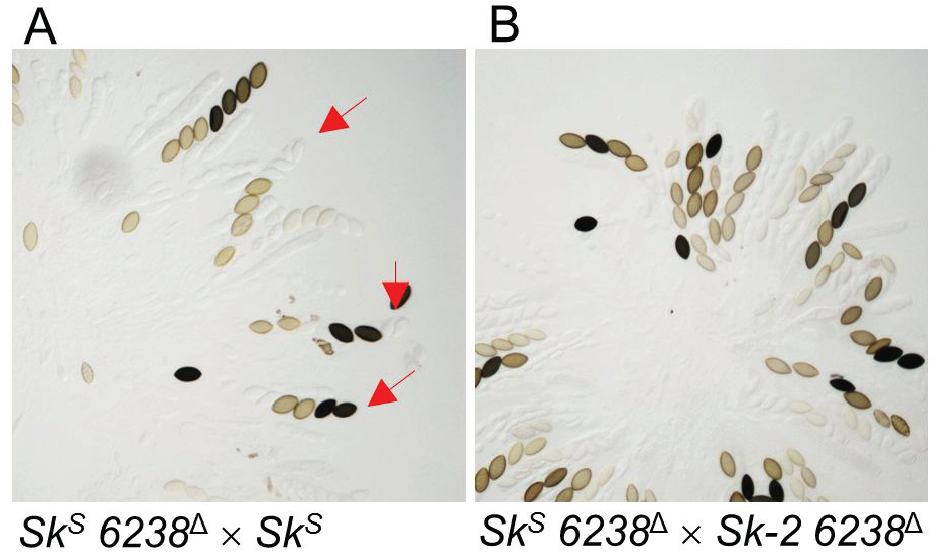
The *ncu06238* gene is not required for spore killing. (A) Asci from a cross between two *Sk*^*S*^ strains (ISU-4559 × P8-43), where one of the strains has had its *ncu06238* coding sequences deleted. While some normal asci are detected, many asci are abnormal and some mimic the spore killer phenotype (red arrows). (B) Asci from a cross between an *Sk*^*S*^ strain and an *Sk-2* strain (ISU-4559 × ISU-4561), where both strains have been deleted of their *ncu06238* coding sequences. Nearly all viable progeny isolated from this cross have the *Sk-2* genotype (47 out of 48, data not shown).

### Replacement of *AH36*^3211^ with *AH36*^*Sk-2*^ restores spore killing to an *rfk-1* mutant

The *rfk-1* mutant strain ISU-3211 carries seven mutations within its *AH36* interval (Figure 6). To confirm that at least one of these mutations (presumably G28326A) is responsible for ISU-3211’s inability to kill ascospores, we replaced *AH36*^3211^ in a descendant of ISU-3211 (strain ISU-3222) with *AH36*^*Sk-2*^::*hph* (Figure 11A and Table S4). Because the presence of an *hph* marker to the right of *AH36* disrupts spore killing in an MSUD-dependent manner (Figure 9B), we performed our test crosses with both a standard *Sk*^*S*^ mating partner and an *Sk*^*S*^ *v150*^Δ^::*hph* mating partner. As expected, we found that replacing *AH36*^3211^ with *AH36*^*Sk-2*^ restores spore killing to a spore killing-deficient strain (Figure 11, B–G). These results demonstrate that the *AH36*^3211^ interval is responsible for the loss of spore killing in ISU-3211 and its *rfk-1* descendants.

**Figure 11.**
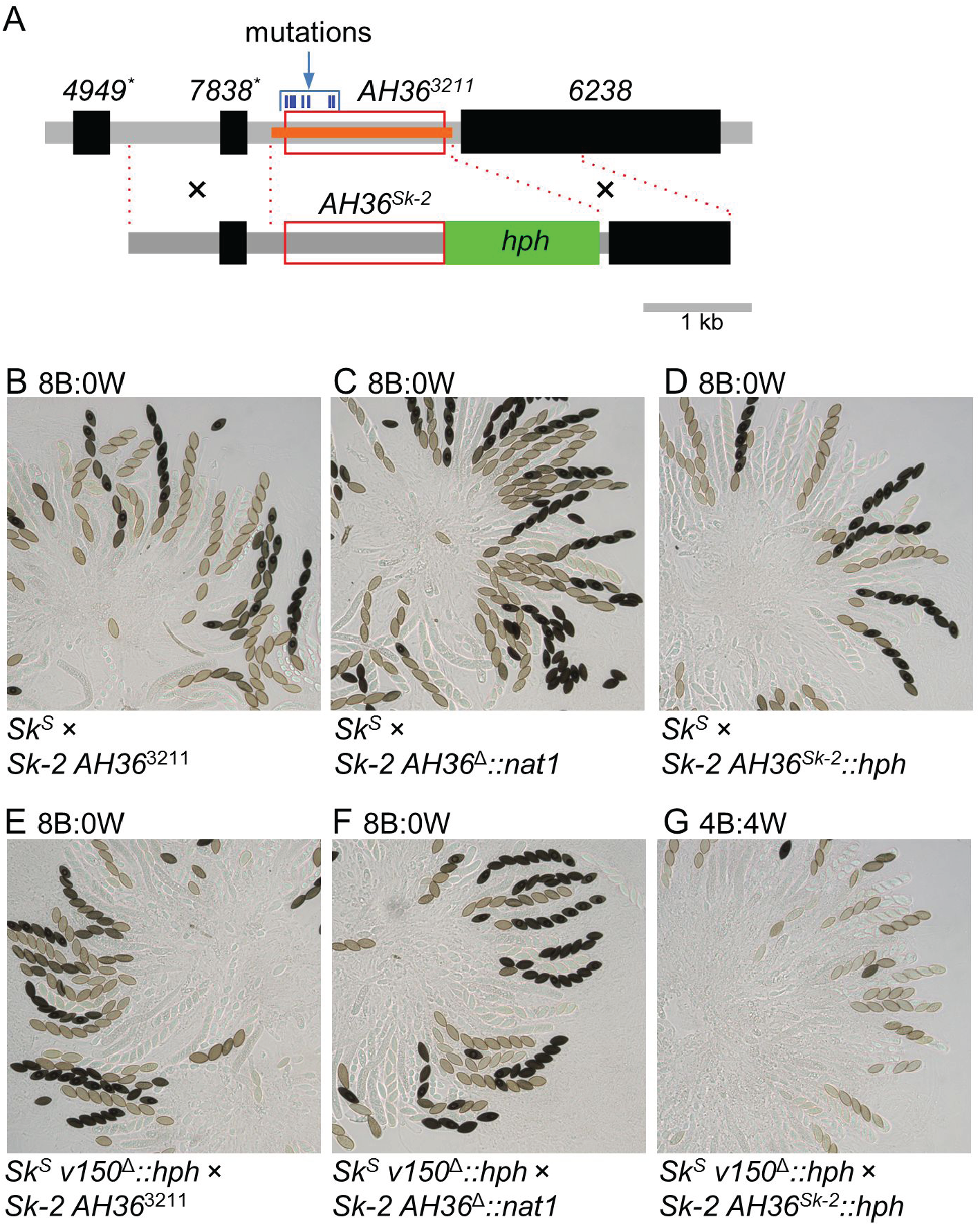
Replacement of *AH36*^3211^ with *AH36*^*Sk-2*^ restores spore killing to an *rfk-1* mutant. (A) Strain ISU-4526 was constructed by replacing *AH36*^3211^ in ISU-3222 (upper diagram, red box) with *nat1* (using vector v160; ISU-3222 is a descendant of ISU-3211). Strain ISU-4563 was then constructed by replacing *AH36*^Δ^::*nat1* in ISU-4562 with *AH36*^*Sk-2*^::*hph* (lower diagram, red box and green rectangle). The locations of the two recombination flanks used to replace *AH36*^Δ^::*nat1* with *AH36*^*Sk-2*^::*hph* are indicated with black crosses and red-dotted lines. (B–D) Asci are from crosses between F2-23 and (B) ISU-3222, (C) ISU-4562, and (D) ISU-4563. (E–G) Asci are from crosses between ISU-4348 and (E) ISU-3222, (F) ISU-4562, and (G) ISU-4563.

### The RFK-1 protein contains (at least) 39 amino acids

Assuming that the start codon for RFK-1 begins at position 28,264, and that the pre-mRNA for *rfk-1* includes no introns (see discussion), we can propose the following hypothesis: RFK-1 is a 39 amino acid protein encoded by DNA located between positions 28,263 and 28,384 (Table 2 and Figure 12A). We found support for this hypothesis by sequencing the *AH36* intervals in strains ISU-3211 through ISU-3216 (Figure 12A), which are the six *Sk-2 rfk-1* isolates obtained by our initial screen for spore killing-deficient mutants (Harvey *et al.* 2014). Specifically, we found that *AH36*^3211^ contains the previously discussed G28326A mutation, which changes the 21^st^ codon from a tryptophan codon to a stop codon; *AH36*^3212^ contains an extra thymine within a run of six thymines between positions 28,281 and 28,288, which causes a frameshift mutation in the 9^th^ *rfk-1* codon; and *AH36*^3213^ contains a G28348A mutation, which changes the 29^th^ codon from an alanine codon to a threonine codon. In addition, we found that the sequences of *AH36*^3214^, *AH36*^3215^, and *AH36*^3216^, are all identical to the sequence of *AH36*^3211^, suggesting that ISU-3211, ISU-3214, ISU-3215, and ISU-3216 were all “fathered” by the same mutagenized conidium. In all, we identified at least one potential codon-altering mutation between positions 28,263 and 28,384 in each of the six known *rfk-1* mutants. This strongly suggests that the interval between positions 28,263 and 28,384 contains at least part, if not all, of the RFK-1 coding sequence.

**Figure 12.**
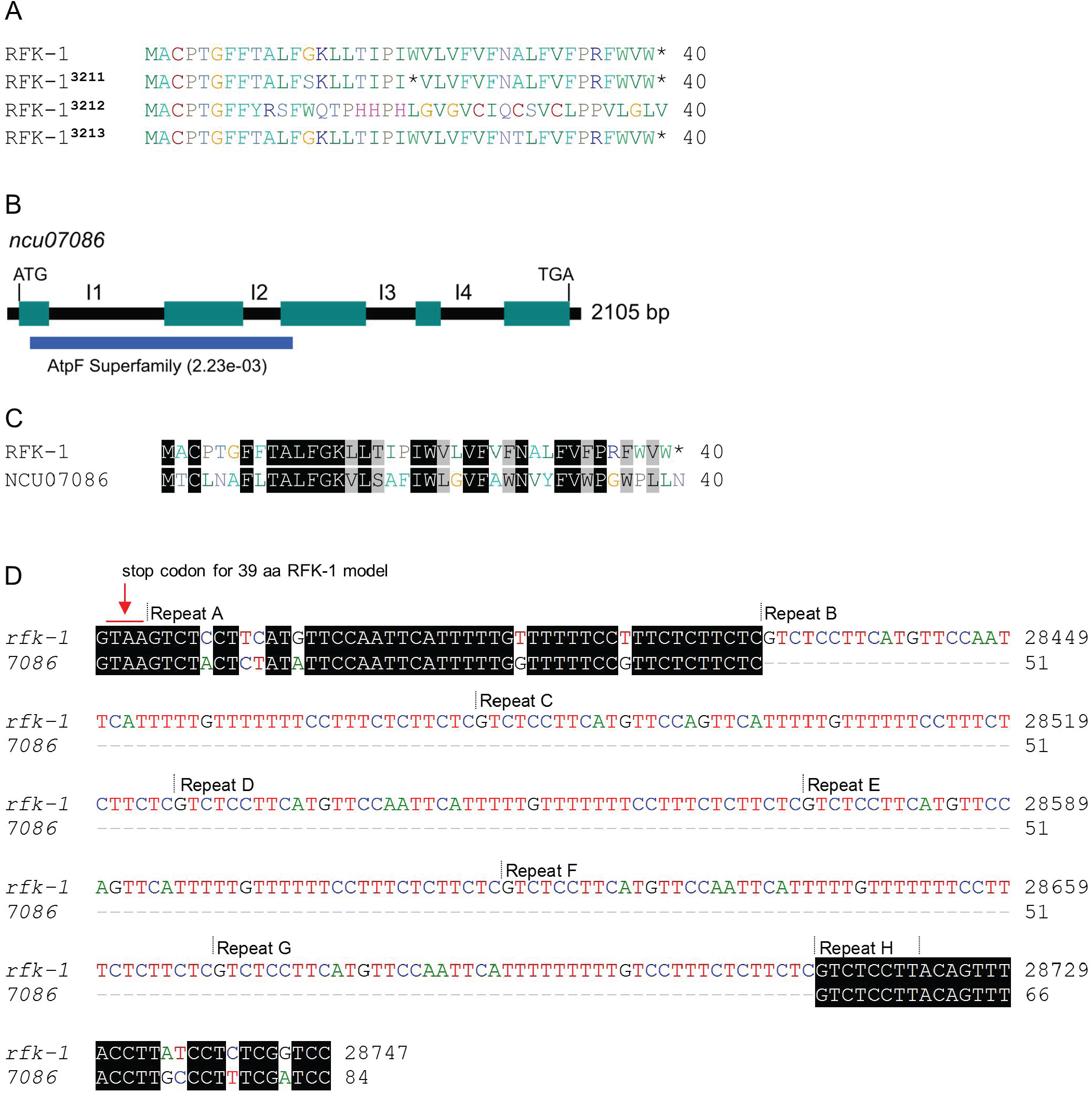
RFK-1 is related to NCU07086. (A) All known *rfk-1* mutations alter the predicted amino acid sequence of RFK-1. (B) A coding region of 2105 bp is predicted for gene *ncu07086* (from start codon to stop codon, including introns). Predicted introns are labeled I1 through I4. An AtpF Superfamily domain (NCBI CDD, Accession No. cl28522) can be identified within the N-terminal end of the NCU07086 protein sequence. (C) The first 39 amino acids of RFK-1 and NCU07086 are similar. (D) The repetitive sequence identified within *AH36* (see Figure 6) appears to have originated from within the first intron of *ncu07086*. The first intron of ncu07086 is predicted to be 141 nucleotides long (NCBI Gene ID: 3876500), and only the first 84 nucleotides are shown. In contrast, the entire *AH36* repeat region is shown along with an additional four nucleotides upstream of “repeat A”. These four nucleotides include the stop codon for the hypothetical 39 amino acid version of RFK-1.

### RFK-1 is related to NCU07086

To investigate the origin of *rfk-1*, we downloaded a list of predicted *N. crassa* proteins from the National Center for Biotechnology Information (NCBI)’s Genome Database (Accession No. GCA_000182925.2) and performed a BLASTP search (Camacho *et al.* 2009) on the list with the hypothetical 39 aa RFK-1 sequence as query (Figure 12A). We found that the most significant match (Expect = 2e-7) to RFK-1 is a hypothetical 362-aa protein called NCU07086 (NCBI Protein Database: XP_960351.1). NCU07086 is encoded by the *ncu07086* gene on *N. crassa* chromosome VI and is predicted to contain four introns (Figure 12B, I1 through I4; NCBI Gene Database, 3876500). A search of NCBI’s conserved domain database (CDD v3.16; Marchler-Bauer *et al.* 2015) with the predicted sequence of NCU07086 identified a region with a low-scoring match to the AtpF Superfamily (Expect=2.32e-3; Figure 12B). Interestingly, RFK-1 is highly similar to the first 39 amino acids of NCU07086 (Figure 12C), and it appears that the 46–48 bp repeat within *AH36* (Figure 6) expanded from a single 47 bp sequence within *ncu07086*’s first intron (Figure 12D). These findings suggest that *rfk-1* evolved from a partial duplication of the *ncu07086* gene.

## DISCUSSION

The biological mechanism used by the Neurospora Spore killers to achieve biased transmission is believed to require the action of a resistance protein and a killer protein. In a previous work, we isolated six *rfk* mutants (ISU-3211 through ISU-3216) and provided evidence that each is mutated at the same locus, subsequently named *rfk-1* (Harvey *et al.* 2014). The *rfk-1* locus in ISU-3211 was mapped to a 45 kb region of *Sk-2*. We began this study with the goal of identifying *rfk-1*. At first, we intended to use three point crossing assays to further refine the position of *rfk-1* within the 45 kb *rfk-1* region. These assays were to be performed with *hph* markers inserted between genes *ncu06192* and *ncu06191* (with vector *v3*) and between genes *ncu06239* and *ncu06240* (with vector *v4*); therefore, deletion vectors *v3* and *v4* were designed to delete relatively small intervals from the *rfk-1* region (25 bp and 261 bp, respectively; Table 2) and they were not expected to influence spore killing. Accordingly, they had no effect on spore killing (Figure 1, C and D). In contrast, *v5* was designed to delete a 10,718 bp interval, spanning most of the *Sk-2*^INS1^ sequence, in hopes that *rfk-1* would be found somewhere within it (Table 2). Fortunately, deletion of interval *v5* (intervals are named after the deletion vectors designed to delete them) was successful and its removal from *Sk-2* eliminated *Sk-2*’s ability to kill ascospores (Figure 1E). We were thus able to focus our efforts on deleting subintervals of *v5,* which allowed us to track *rfk-1* to the intergenic region between *ncu07238\** and *ncu06238* (Figures 2, 3, and 10).

We also tested various subintervals of *v5* for the presence of *rfk-1* by transferring them to an *Sk*^*S*^ strain and performing test crosses with an *Sk*^*S*^ *sad-2*^Δ^ mating partner (Figure 4). For this assay to yield positive results, *rfk-1* must be sufficient for spore killing. Indeed, we found this to be the case when we identified four intervals (*AH30, AH31, AH37*, and *AH36*) that trigger ascus abortion. These four intervals all have the 1481 bp of *AH36* in common, and the ascus abortion phenotype associated with each interval is likely due to the presence of *rfk-1* without a compatible resistance gene. For example, the KN model holds that the resistance protein (RSK) and the killer are both active during early stages of meiosis (Hammond *et al.* 2012). Lack of a resistant version of RSK, along with expression of the killer, may cause asci to abort meiosis before ascospore delimitation. This phenomenon explains the abortion phenotypes of *AH30, AH31,* and *AH37*. However, for succinctness, we also referred to the phenotype associated with *AH36* as ascus abortion, although it may be more accurate to refer to it as a “bubble” phenotype. The bubble phenotype was originally described by Raju *et al*. (1987), and it is thought to arise when asci and/or ascospores abort shortly after ascospore delimitation. Therefore, one explanation for the existence of the two phenotypic classes is that ascus development progresses a bit further with *AH36* than it does with *AH30, AH31,* and *AH37.* Asci could progress further with *AH36* if *rfk-1* expression is lower from *AH36* than it is from *AH30, AH31,* and *AH37*. In line with this reasoning, *AH36* is the shortest of the abortion-inducing intervals, and, as a result, it may lack some of the regulatory sequences needed for full expression of *rfk-1*. It should be possible to address this hypothesis once the complete transcriptional unit of *rfk-1* is identified.

Although we have yet to identify *rfk-1*’s transcriptional start (+1) site and termination site, or confirm the presence/absence of introns, we have provided strong evidence that the *rfk-1* coding region includes the DNA interval between positions 28,263 and 28,384 (Figure 6). For example, a putative nonsense mutation at position 28,326 disrupts the ascus-aborting ability of interval *AH36* (Figure 7); spore killing functions when a non-native promoter is attached to the putative RFK-1 start codon at position 28,264 (Figure 8); all six of the known *rfk-1* mutants carry putative codon-altering mutations between positions 28,263 and 28,384 (Figure 12A), and insertion of a non-native promoter in the middle of this region disrupts spore killing (Figure 8). However, while our data indicate that the positions between 28,263 and 28,384 are part of the *rfk-1* coding region, they do not eliminate the possibility that the coding sequences for RFK-1 include additional positions upstream and/or downstream of 28,263 and 28,384, respectively. Indeed, our preliminary analysis of RNAseq data from *Sk*^*S*^ *× Sk-2* crosses (unpublished data) strongly suggests that an intron may exist between positions 28,379 and 28,775. The 5’ splice site of this hypothetical intron is related to the 5’ splice site of the first intron of *ncu07086* (Figure 12D). If this intron does exist within the *rfk-1* pre-mRNA, the RFK-1 stop codon would shift downstream and the length of RFK-1 would increase to 101 aa (assuming position 28,264 is the start codon and no other introns influence the stop codon position). Future work will seek to fully characterize the *rfk-1* coding region by identifying the transcriptional start site, termination site, and any introns that may exist for the primary *rfk-1* transcript, as well as for any biologically significant variants, if they were to exist.

While this work represents a significant step towards understanding the mechanism of *Sk-2-*based spore killing, many questions remain unanswered. For example, although it appears that RFK-1 evolved from NCU07086, does RFK-1 interfere with NCU07086 function as part of the spore killing mechanism? NCU07086 contains a region with slight homology to the AtpF Superfamily (Figure 12D). Interestingly, the *atpF* gene in *E. coli* (also known as *uncF*; NCBI Gene ID 948247) encodes subunit b of the F-type ATP synthase complex (Walker *et al.* 1984; Dunn 1992; McLachlin and Dunn 1997; Revington *et al.* 1999). This hints that RFK-1 could mediate spore killing by targeting eukaryotic F-type ATP synthases, which are associated with mitochondrial membranes in eukaryotes (Stewart *et al.* 2014). However, NCU07086 in *N. crassa* has not been investigated and a much more likely candidate for the b subunit of *N. crassa*’s F-type ATP synthase is found in NCU00502 (KEGG oxidative phosphorylation pathway: ncr00190, release 87.0, Kanehisa and Goto 2000; Kanehisa *et al.* 2016). Thus, at this point in time, a role for RFK-1 in disrupting mitochondrial function as part of the spore killing process is purely speculative.

Although the primary goal of this work was to identify *rfk-1*, the identity of which has been of interest to meiotic drive researchers since the discovery of *Sk-2* nearly four decades ago, we unexpectedly discovered the strongest evidence to date that genomes in some, if not all, lineages of eukaryotic organisms possess elaborate defense processes to protect themselves from meiotic drive. With respect to Neurospora genomes, this defense process appears to be MSUD. The first hint that MSUD defends Neurospora genomes from meiotic drive appeared in 2007, when it was discovered that *Sk-2* and *Sk-3* are weak MSUD suppressors (Raju *et al.* 2007). Next, in 2012, it was found that the position of *rsk* within *Sk-2* allows it to pair with *rsk* in the *Sk*^*S*^ genome during *Sk-2* × *Sk*^*S*^ crosses. If *rsk* is not paired during these crosses (*e.g.*, if it is deleted from the *Sk*^*S*^ mating partner), it is silenced by MSUD and the entire ascus is killed by the killer protein, which we now know to be RFK-1. In the current work, we found that the position of *rfk-1* within *Sk-2* is also critical for the success of meiotic drive because it allows *rfk-1* to escape inactivation by MSUD. However, unlike *rsk, rfk-1* is only found in *Sk-2* strains and it cannot be paired in *Sk-2* × *Sk*^*S*^ crosses. Evolution appears to have found a way to circumvent this problem by positioning *rfk-1* close to sequences that are paired during meiosis (*i.e.* close to *ncu06238* in Figure 10). Our data indicate that the proximity of *rfk-1* to paired sequences allows it to escape inactivation by MSUD, which is critical for the success of spore killing. Overall, our findings add to accumulating evidence that MSUD antagonizes the evolution of meiotic drive elements by placing significant constraints on the arrangement of critical genes within the elements. Furthermore, our findings suggest that eukaryotic genomes like those of Neurospora fungi have evolved elaborate defense mechanisms to protect themselves from meiotic drive.

## ACKNOWLEDGEMENTS

We are grateful to members of the Brown, Hammond, Johannesson, and Shiu laboratories for assistance with various technical aspects of this work. We are pleased to acknowledge use of materials generated by P01 GM068087 “Functional Analysis of a Model Filamentous Fungus”. We are also grateful to the Fungal Genetics Stock Center, whose preservation and distribution of Neurospora isolates helped make this work possible (FGSC; McCluskey *et al*. 2010). This project was supported by a grant from the National Science Foundation (MCB# 1615626) (T.M.H.). P.K.T.S was supported by the National Science Foundation (MCB# 1715534) and the University of Missouri Research Board and Research Council. H.J. was supported by the European Research Council grant under the program H2020, ERC-2014-CoG, project 648143, and the Swedish Research Council. Mention of trade names or commercial products in this article is solely for the purpose of providing specific information and does not imply recommendation or endorsement by the U.S. Department of Agriculture. USDA is an equal opportunity provider and employer.

## Supporting Information

Rhoades *et al*. “Identification of a genetic element required for spore killing in Neurospora”.

**Figure S1.**
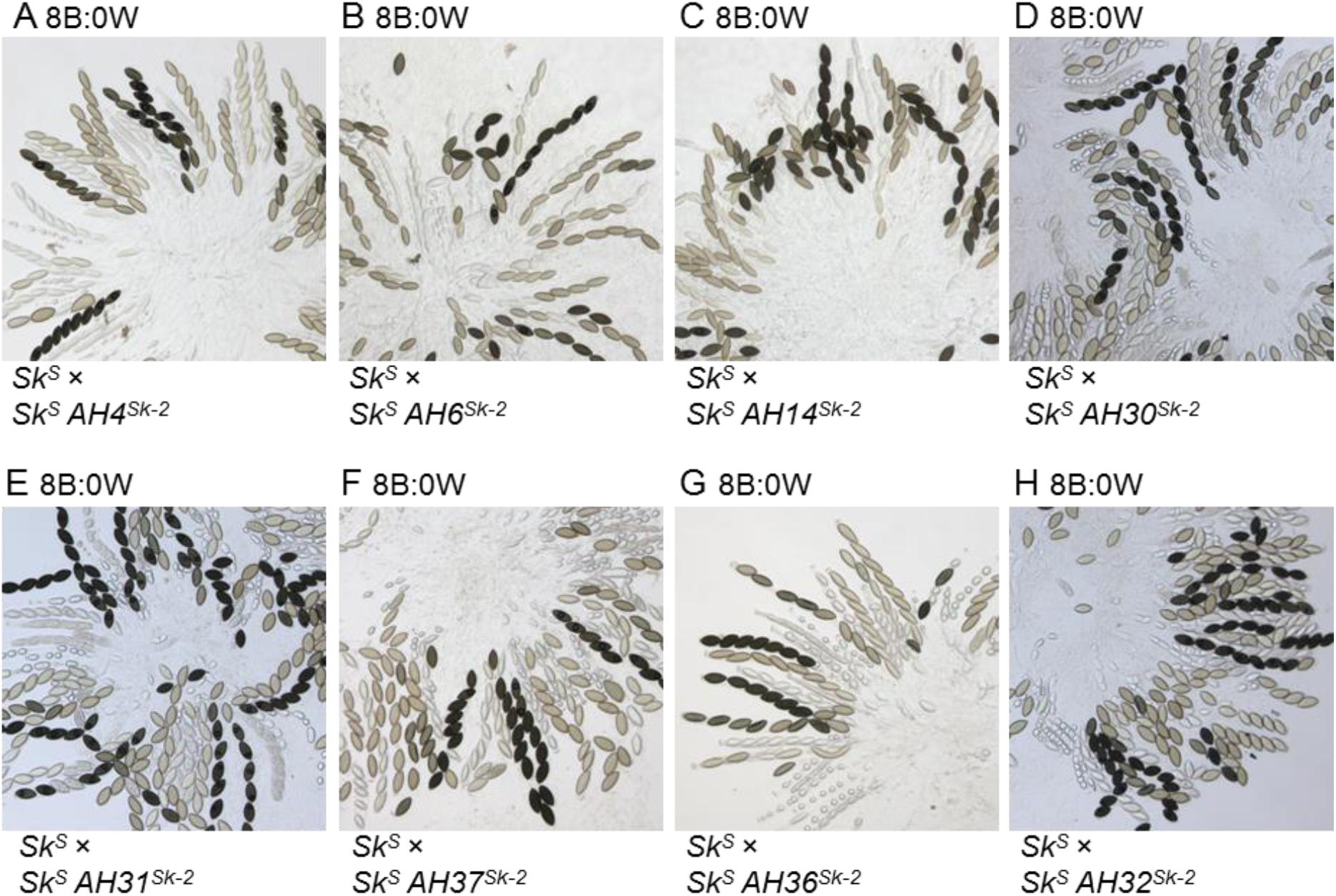
Unpaired *Sk-2*^INS1^-intervals do not kill ascospores in MSUD-proficient crosses. Unpaired *Sk-2*^INS1^-intervals do not kill ascospores in MSUD-proficient crosses. (A–H) The images depict asci from crosses between *Sk*^*S*^ strains, one of which carries an interval of the *Sk-2*^INS1^ locus (*e.g., AH4*^*Sk-^2^*^, *AH6*^*Sk-^2^*^, etc.). All crosses produced asci with an 8B:0W phenotype. Crosses are as follows: (A) F2-26 × ISU-3224, (B) F2-26 × ISU-3228, (C) F2-23 × ISU-3243, (D) F2-26 × ISU-3656, (E) F2-26 × ISU-3658, (F) F2-23 × ISU-4269, (G) F2-26 × ISU-4271, and (H) F2-26 × ISU-3660.

**Table S1.**
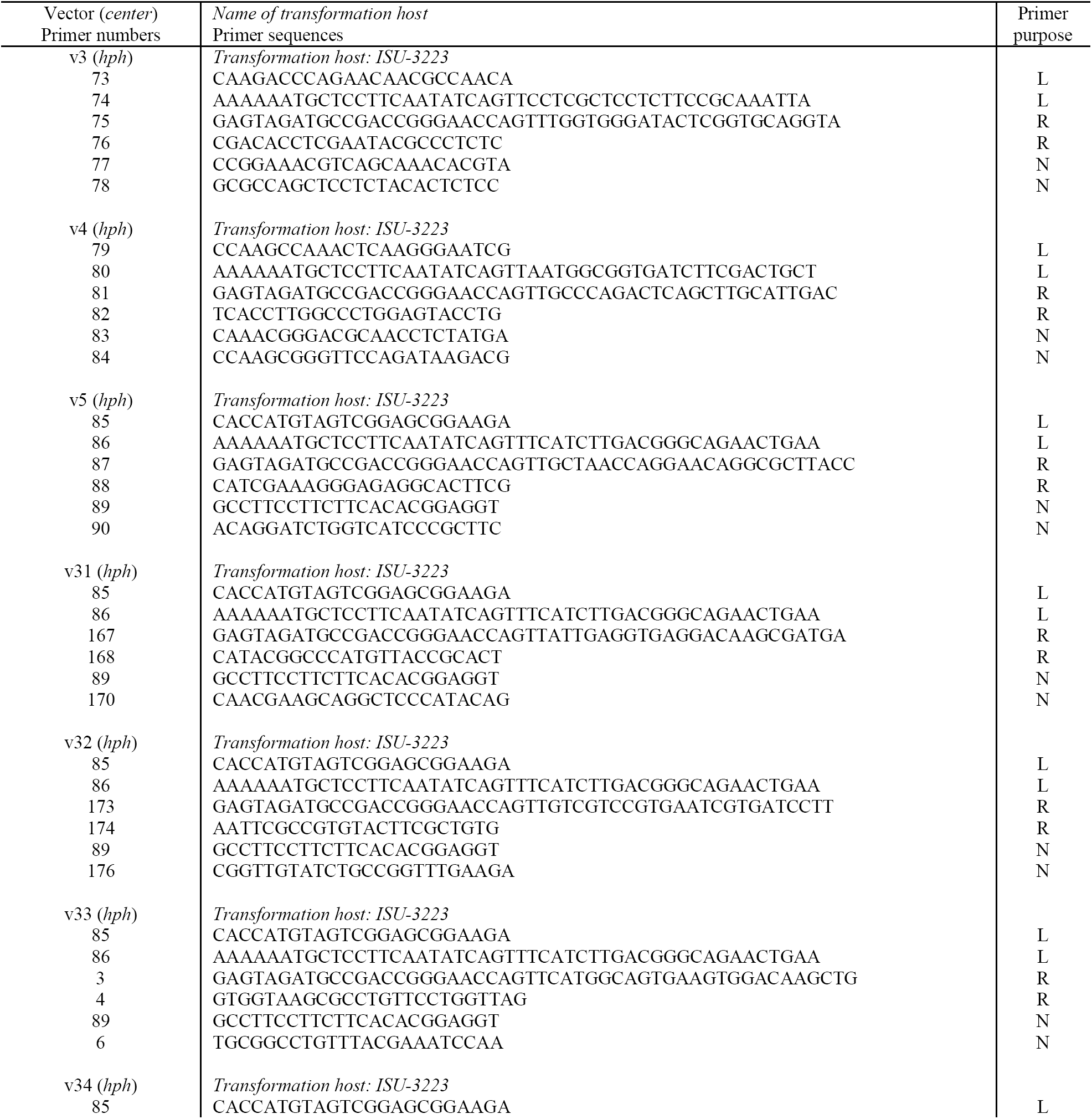
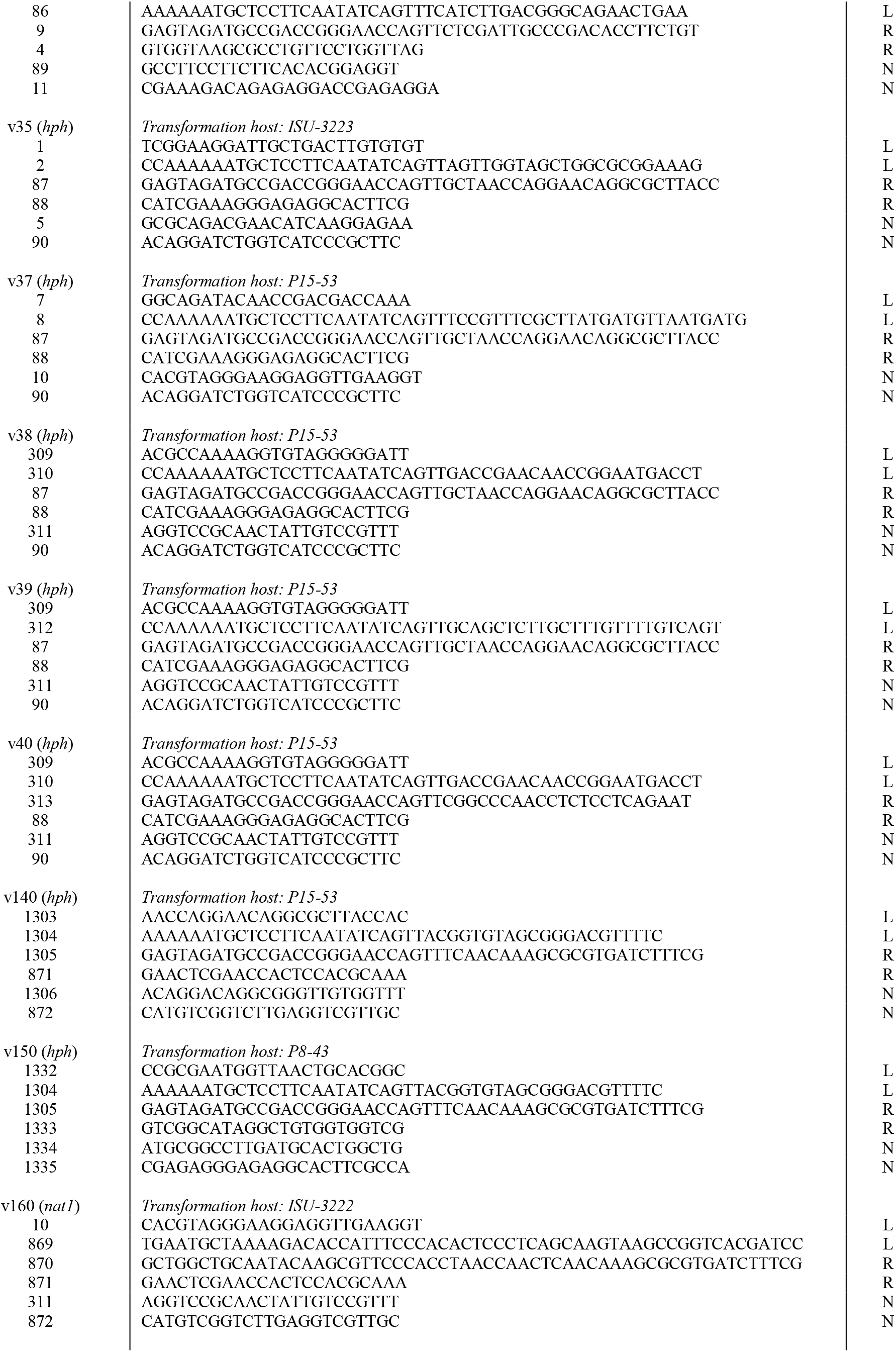
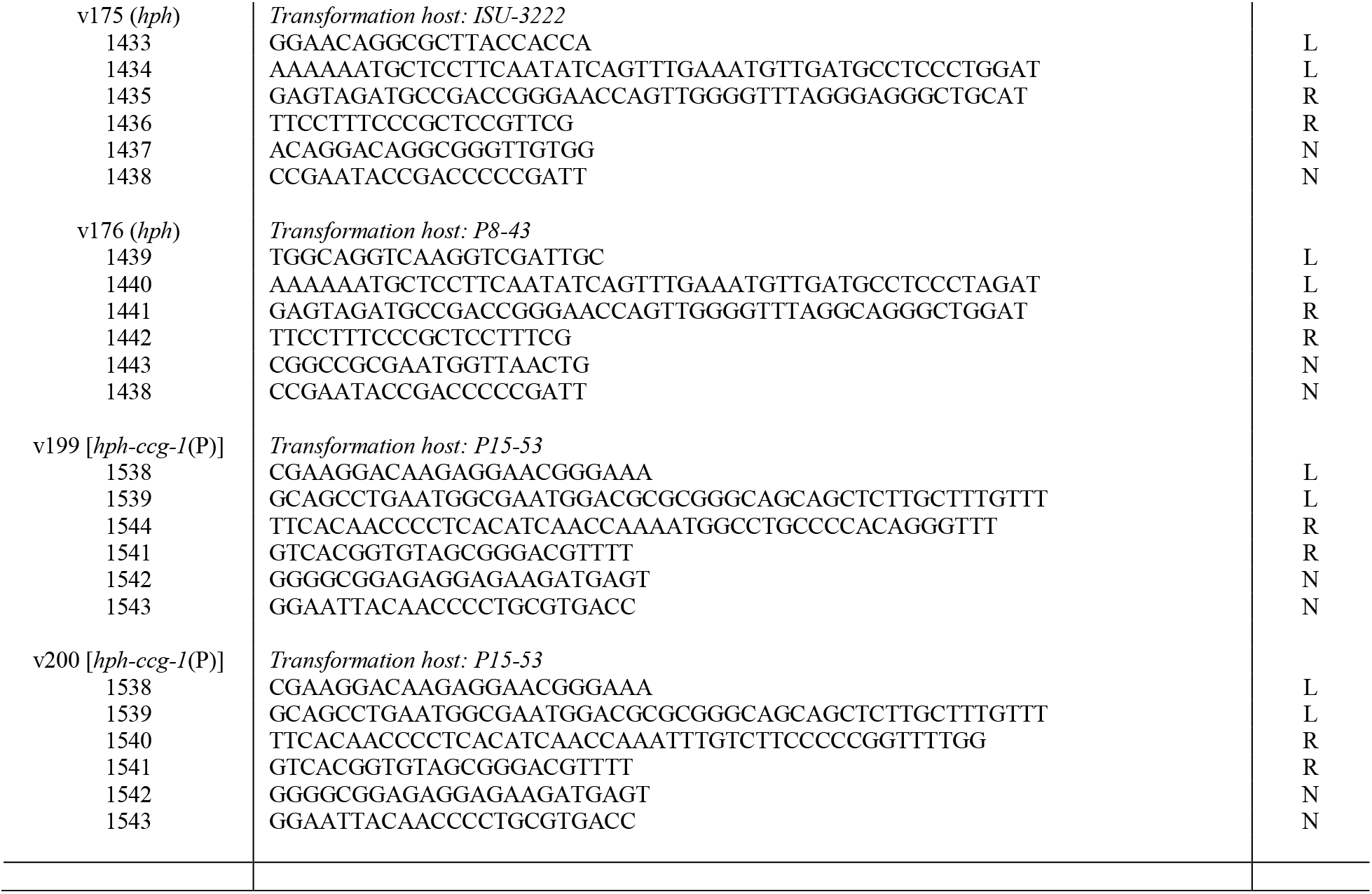
Primers for DJ-PCR-based construction of deletion vectors. Nineteen deletion vectors were constructed by double joint (DJ)-PCR (Yu *et al.* 2004; Hammond *et al.* 2011). The table below lists the forward and reverse primer sequences (5′ to 3′) for the left recombination flank (L), the right recombination flank (R), and the nested amplification of each completed vector (N). For each vector, the left and right DNA flanks were amplified from genomic DNA of the transformation host, which is also indicated in the table. The center fragment for each vector is listed next to the name of each vector in the left-most column. Center fragments were either *hph, nat1*, or *hph-ccg-1*(P). See Table S2 for more information on the center fragments.

**Table S2.**
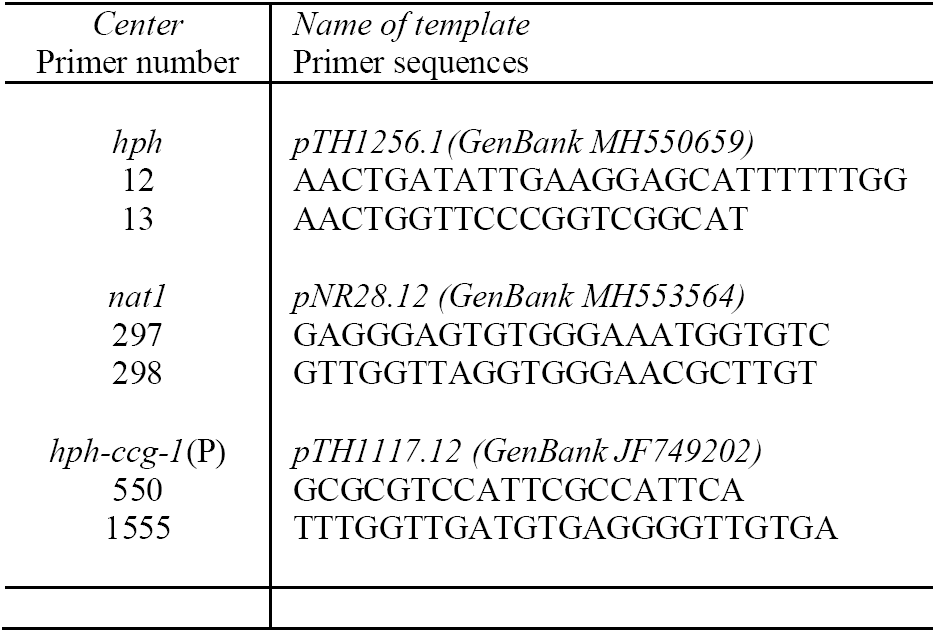
Primers for DJ-PCR center products. The forward and reverse primers used to amplify the center fragments for construction of DJ-PCR deletion vectors are described below.

**Table S3.**
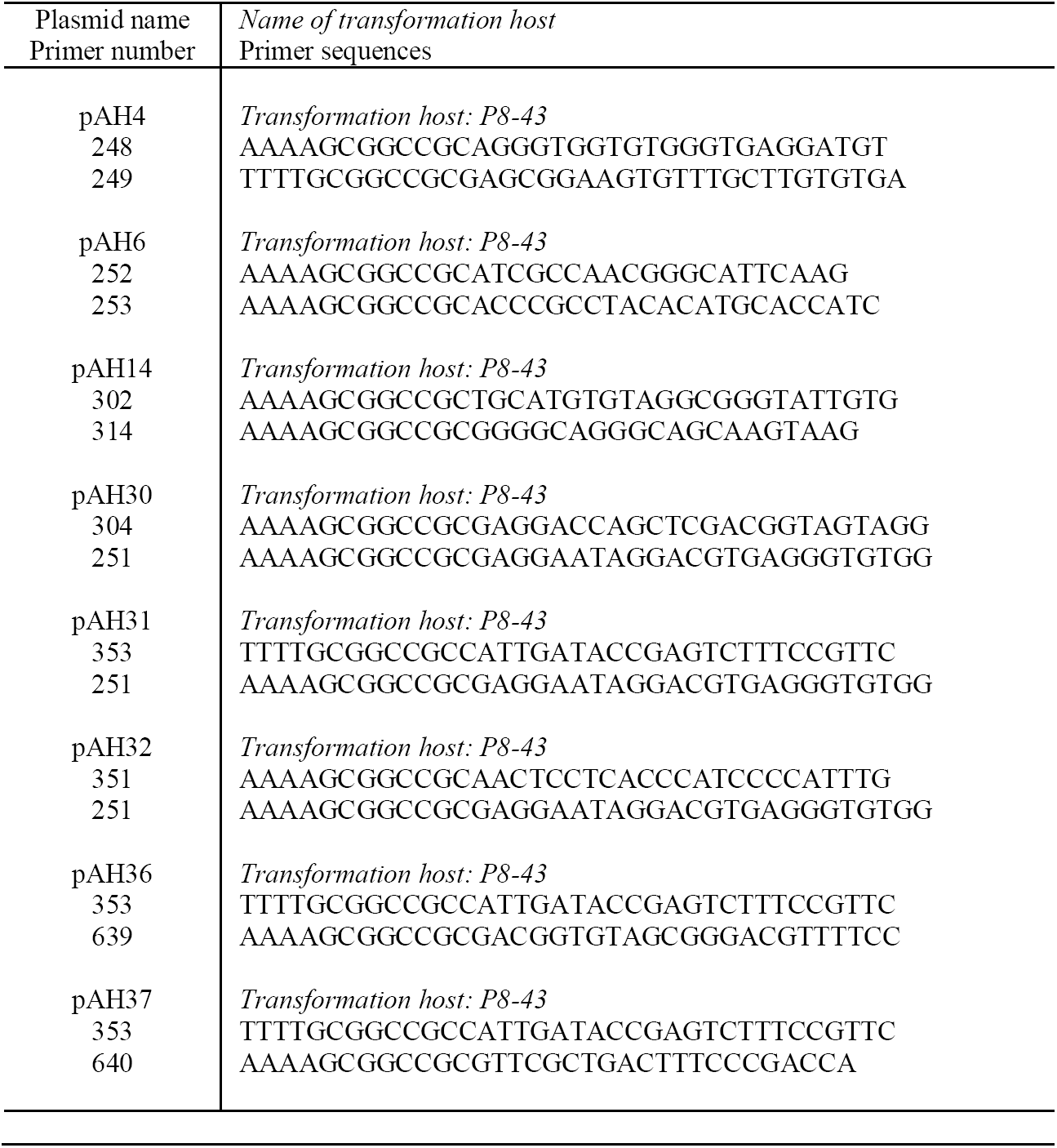
Primers for cloning *Sk-2* intervals to pTH1256.1. Eight intervals of *Sk-2*^INS1^ were cloned to the *Not*I site of pTH1256.1 (GenBank MH550659), using the primers listed below. These cloning procedures created plasmids pAH4, pAH6, pAH14, pAH30, pAH31, pAH32, pAH36, and pAH37. Each plasmid was then used to transform strain P8-43.

**Table S4.**
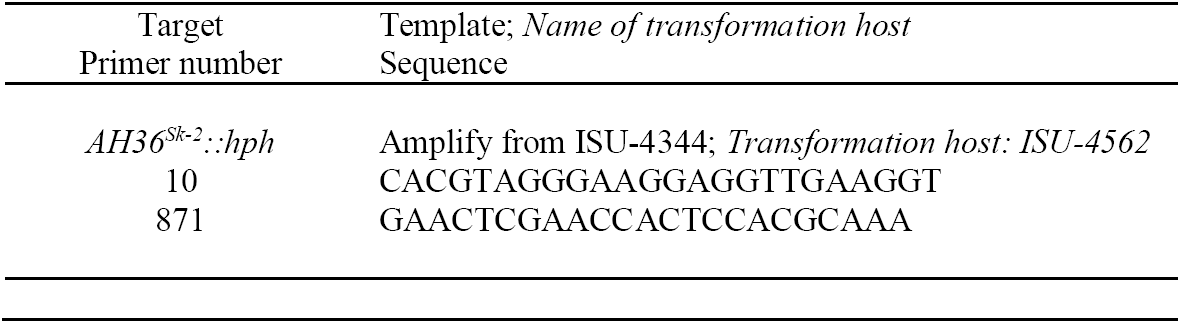
Primers for amplification of *AH36*^*Sk-^2^*^::*hph.* The *AH36*^*Sk-^2^*^::*hph* allele was amplified from ISU-4344 using the primers 10 and 871. These primers span the *v140*^Δ^::*hph* allele in ISU-4344 and produce a PCR product containing *AH36*^*Sk-^2^*^ and *hph* between recombination flanks suitable for replacing *AH36*^Δ^::*nat1* in ISU-4562 with *AH36*^*Sk-^2^*^::*hph*.

**Table S5.**
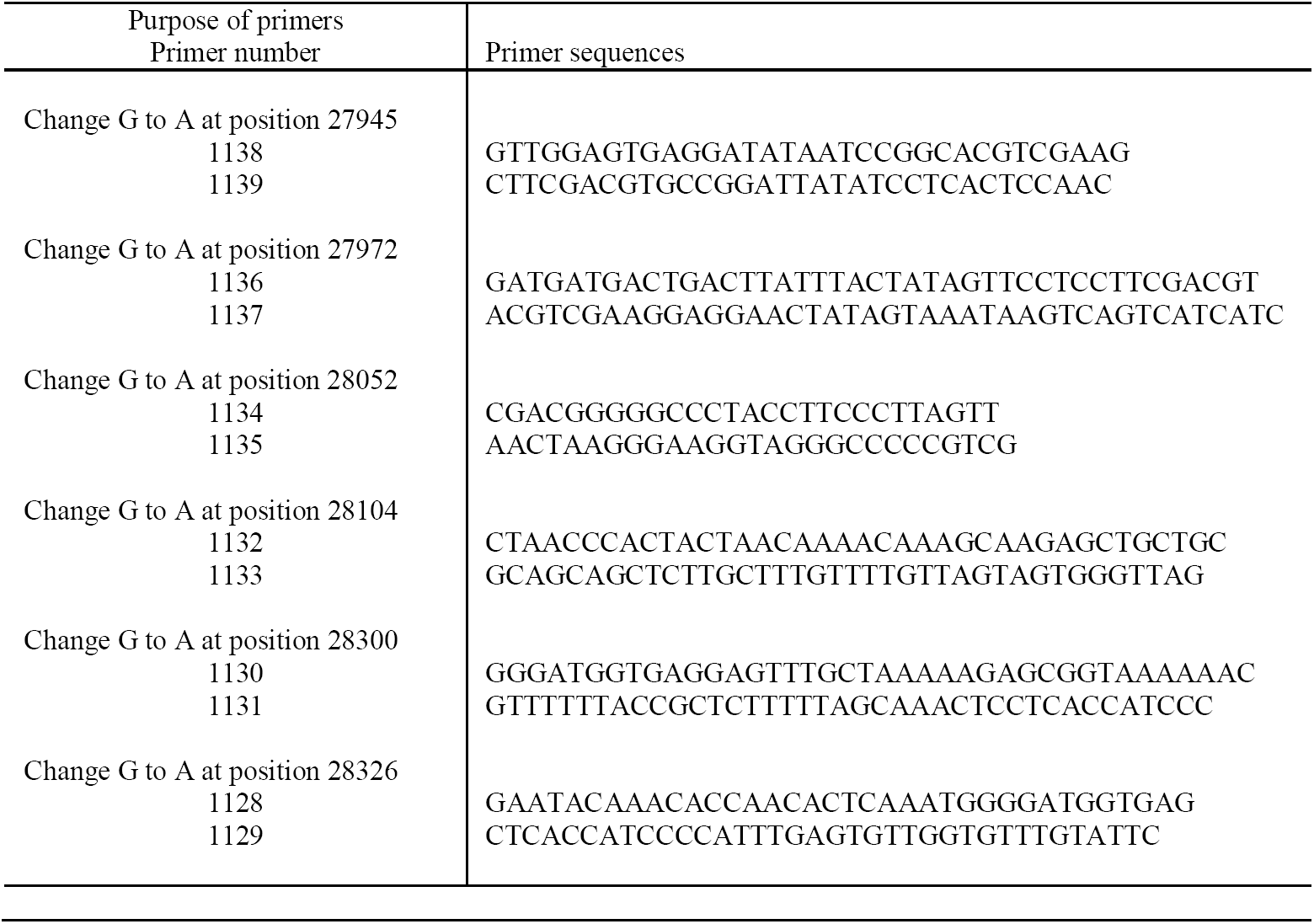
Primers for site directed mutagenesis of *AH36*^*Sk-^2^*^. Site directed mutagenesis was performed essentially as described for the QuikChange II Site-Directed Mutagenesis Kit (Revision E.01, Agilent Technologies). The *AH36* interval from *Sk-2* was cloned to the *Not*I site of a standard 3 kb bacterial cloning vector with primers 353 and 639 (Table S2). Site-specific mutations were introduced into the resulting plasmid (pNR9.1) by PCR with the primer sets described below. PCR products were digested with *Dpn*I and used to transform chemically-competent *E. coli* Ig™ 5-alpha cells (Intact Genomics). Site directed mutations were confirmed by Sanger sequencing and mutated plasmids were used to transform P8-43.

